# Diverse microbial metal resistance and novel metal cycling organisms in copper/nickel mine tailings

**DOI:** 10.64898/2026.02.27.708553

**Authors:** Molly Chen, Daniel S. Grégoire, Jeffrey G. Bain, David W. Blowes, Laura A. Hug

## Abstract

Mine tailings contribute to environmental heavy metal contamination through the formation of acid mine drainage (AMD). Microbially-mediated processes such as iron and sulfur redox cycling influence metal mobility. Here, we applied an integrated metagenomic and metaproteomic approach to profile microbial communities across vertical geochemical gradients in legacy copper/nickel tailings in Sudbury, Ontario, Canada. From 43 samples, we recovered 454 non-redundant metagenome-assembled genomes (MAGs), revealing diverse populations within the Actinobacteriota, Desulfobacterota, and uncultured lineages such as *Candidatus* Eremiobacterota and SZUA-79. Functional profiling identified 301 putative iron- and sulfur-cycling MAGs, including those within the *Ca.* Eremiobacterota and SZUA-79 phyla. Metal resistance genes were widespread and diverse, with abundances that did not correlate with measured metal concentrations. Proteomic data confirmed *in situ* expression of selected metal resistance genes and iron/sulfur metabolism genes, despite limited protein recovery from this challenging matrix. Our findings highlight both the depth of microbial diversity in metal resistance and metal biogeochemical cycling in mining waste, as well as the technical challenges that currently limit genomic and proteomic sequencing coverage in low-biomass, metal-rich matrices. This work also presents new protocols for multi-omics data capture and analysis from metal contaminated environments, including new protein extraction and bioinformatic gene annotation tools.

**Highlights:**

- Metagenomics recovered 454 non-redundant MAGs from copper/nickel mine tailings, revealing high taxonomic novelty including uncultured *Ca.* Eremiobacterota and SZUA-79 lineages.
- A custom HMM-based pipeline targeting metal resistance genes uncovered a widespread and diverse set of metal resistance genes across MAGs in tailings, which did not correlate with metal concentrations.
- Metaproteomics validated *in situ* expression of both iron/sulfur cycling genes and metal resistance proteins, although low biomass and contamination limited proteomic sequencing coverage.

## Introduction

Mine tailings are waste materials generated from the extraction of valuable metals from ores. After crushing ores into smaller particles and separating metals and minerals via flotation, density, and leaching (Blowes et al., 2003), the residual fine-grained minerals and waste rock are typically mixed with water, forming a high-volume slurry of tailings that is transported to a designated storage facility (Blowes et al., 2003).

Sulfide-rich tailings pose an on-going risk of environmental contamination due to the oxidation of sulfide minerals such as pyrite (FeS_2_), which leads to the formation of Acid Mine Drainage (AMD). AMD is characterized by moderately (pH<7) to extremely (pH<3) acidic wastewater with high concentrations of heavy metals (Johnson & Hallberg, 2003). Although mineral oxidation can occur abiotically, the activity of acidophilic chemolithoautotrophs (*e.g., Acidithiobacillus ferrooxidans*, *Leptospirillum ferrooxidans*, *Sulfolobus metallicus*) can significantly accelerate the rate of AMD formation by oxidizing ferrous iron (Fe^2+^) to ferric iron (Fe^3+^) and reduced inorganic sulfur compounds (RISCs) to sulfate (SO_4_^2-^) (Johnson & Hallberg, 2003; Newsome & Falagán, 2021). These processes release AMD into soils and freshwater ecosystems, which highly detrimental to environmental and human health (reviewed in Simate & Ndlovu, 2014).

Mining waste microbial communities must possess adaptations to overcome the challenge of living in a harsh environment characterized by low pH and high concentrations of dissolved heavy metals. These adaptations rely on the use, acquisition, and/or evolutionary adaptation of metal resistance genes (MRGs). MRGs include membrane transporters for exporting metal ions outside of the cell (Argüello et al., 2007), oxidoreductases for changing the redox state of metal ions into less toxic or bioavailable forms (*e.g.,* from Cu^2+^ to Cu^1+^) (Grass & Rensing, 2001), and metal-binding proteins for sequestering metal ions in the cytoplasm or periplasm (Argüello et al., 2013). Additionally, many microbes inhabiting tailings utilize iron- and sulfur-cycling genes in energy-generating metabolic pathways: these include iron- and sulfur oxidizing chemolithoautotrophs (examples above), as well as iron- and sulfate-reducing organisms (*e.g.*, *Acidiphilium* spp., *Desulfovibrio* spp.) that use ferric iron and/or sulfate as alternative electron acceptors in anoxic tailings environments (Johnson & Hallberg, 2003); such genes will be referred to in this study as metal metabolism genes, or MMGs.

Understanding the microbial activities that control metal biogeochemical cycling in tailings is crucial for predicting the fate of metals in these systems and developing effective remediation strategies. For example, cover layers installed above tailings can restrict oxygen ingress, limit oxidation and facilitate anaerobic microbial metabolisms such as sulfate reduction, promoting secondary sulfide mineral precipitation (Johnson & Hallberg, 2005). To target specific microbial metal-cycling activities, a comprehensive profile of microbial communities, including the MRG/MMGs they express, is required.

A previous investigation into tailings microbial communities identified high variability in community composition and diversity along depth gradients, and across four legacy copper/nickel tailings sites in Sudbury, Ontario (Chen et al., 2024). The extent of sulfide mineral oxidation and presence of remedial cover layer(s) varied between sampling locations, influencing the relative acidity and dissolved metal concentrations of the environments. Through 16S rRNA amplicon sequencing on 10 cm vertical sections of tailings cores, we determined a high-resolution taxonomic profile of communities across a range of geochemical gradients. This study identified correlations between microbial community structures with pH, Eh, alkalinity, salinity, and some metal ions, which unexpectedly did not include Cu or Ni. Many lineages associated with iron/sulfur cycling in tailings were identified, as well as a number of novel taxa that have not previously been described or studied within mine tailings (Chen et al., 2024).

Although 16S rRNA amplicon sequencing is a valuable tool for characterizing environmental microbial diversity, it does not allow for direct inference of the potential metabolic activities or functions of organisms. Predicting mechanisms of heavy metal resistance and biogeochemical cycling in mine tailings is improved by the use of metagenomics to identify MRGs and MMGs, especially those that are taxonomically novel or lack isolated representatives. Additionally, the activity and subsequent ecological impact of organisms in their natural environments are not necessarily tied to taxonomic or gene abundances. These roles can be clarified by measuring gene expression (*i.e.*, via RNA and/or proteins) (Wang et al., 2016). Metagenomic, metatranscriptomic, and metaproteomic (‘multi-omics’) studies of AMD environments have been crucial to advancing our knowledge of mining waste microbiology (reviewed in Huang et al., 2016; Méndez-García et al., 2015), providing information about the importance of rare populations (Hua et al., 2015), metal resistance strategies (Ayala-Muñoz et al., 2020; Mirete et al., 2007), ecological succession (Mueller et al., 2011), and potential for biomining (Cárdenas et al., 2016; Kumar et al., 2023). Metaproteomics even enabled the discovery of a novel cytochrome essential for iron oxidation within AMD communities (Ram et al., 2005).

It can be challenging to collect sequencing data from mine tailings due to the low biomass, high volume sample matrix, and the presence of heavy metal contaminants, which can interfere with processes such as DNA recovery and amplification (Bonsu et al., 2024). Currently, there are no commercial products optimized for nucleic acid or protein extraction from tailings as there are for other types of microbiomes (*e.g.*, soil, gut). Metagenomic studies on tailings communities have been conducted (Ghosh & Das, 2018; Gupta et al., 2017; Liu et al., 2021), but metaproteomic studies of mining waste have focused on biofilm samples (reviewed in Wilmes et al., 2015). There have been no proteomic analyses of biomass extracted directly from mine tailings to date, likely due to difficulties extracting from these complex matrices.

The objective of this study was to combine metagenomic and metaproteomic approaches to identify microbial metal resistance mechanisms and iron/sulfur cycling capabilities in sulfidic copper/nickel mine tailings, located in two tailings areas in Sudbury, Ontario, Canada. To meet this objective, we developed a method of protein extraction from mine tailings, a challenging, low-biomass matrix. We used a combination of existing and custom-generated gene annotation tools to characterize MRG and MMGs (including sulfur-cycling genes) present in microorganisms inhabiting mine tailings. In doing so, we identified organisms of interest with high metal resistance and/or metal cycling potential, and considered how differences in MRG and MMGs between populations may be influenced by environmental variables such as metal concentrations.

## Results and discussion

### Sampling site characteristics

Sampling occurred in November 2021 from four sampling locations at two sulfide tailing impoundments operated by Glencore Sudbury Integrated Nickel Operations (Sudbury INO): ML25 and ML34, from the Strathcona Tailings Management Area, which received tailings from 1968 to 2012; as well as NR18 and NR3, from the Nickel Rim North tailings area, which received tailings from 1953 to 1958 (Bain et al., 1998; Bain & Blowes, 2013; Chen et al., 2024; Johnson et al., 2000).

ML25 tailings are highly oxidized and pore-water metal concentrations are relatively high compared to ML34, NR18, and NR3. Tailings near ML34 are overlain by an organic cover layer (biosolids and municipal compost) that was around 0.4 m thick at the time of sampling, limiting oxidation of the underlying sulfide minerals, which was included in the sample material. Additional details on mineral composition, storage conditions, and contaminant profiles for each of the sampling locations are described here when relevant, and more fully in (Chen et al., 2024).

### Sequencing statistics

A total of 757 high-quality (>70% complete, <10% contaminated) metagenome-assembled genomes (MAGs) (454 non-redundant, after dereplication) were identified across 43 samples (ranging from 1–54 per sample, average of 18) after assembly and binning of metagenomic sequencing data (**Table 1**). The number of high-quality MAGs per sample was much lower than the number of ASVs identified through 16S rRNA amplicon sequencing (minimum=141) (Chen et al., 2024). One sample (ML25_013) did not return any high-quality bins and was excluded from subsequent analyses involving MAG gene annotation. Low MAG counts were often associated with a low fraction of assembled reads (**Table 1**), likely resulting from insufficient sequencing coverage of low biomass samples, despite PCR amplification during the metagenome library preparation step.

**Table 1:** Metagenomic sequencing statistics by sample. Scaffolds were binned into metagenome-assembled genomes (MAGs) using MetaBAT2, CONCOCT, and MaxBin2, and DAS Tool was used to identify the highest quality set of non-redundant bins among the three binning algorithms. CheckM was used to filter for bins with >70% completeness and <10% contamination as the final set of high-quality

| Sample | Depth (m) | Reads | % Assembled | Scaffolds | Scaffolds binned | HQ MAGs |
| --- | --- | --- | --- | --- | --- | --- |
| ML25_003 | 0.28 | 16,647,582 | 76.03 | 15,966 | 8,387 | 33 |
| ML25_006 | 0.62 | 21,542,193 | 79.92 | 18,432 | 7,081 | 32 |
| ML25_007 | 0.73 | 13,168,935 | 84.90 | 8,715 | 4,309 | 22 |
| ML25_008 | 0.84 | 16,652,969 | 89.41 | 8,187 | 3,727 | 19 |
| ML25_009 | 0.96 | 36,061,742 | 95.62 | 6,644 | 3,579 | 24 |
| ML25_010 | 1.07 | 37,476,942 | 89.22 | 4,672 | 2,323 | 13 |
| ML25_012 | 1.29 | 3,838,337 | 30.03 | 1,985 | 679 | 2 |
| ML25_013 | 1.42 | 4,011,559 | 0.59 | 7 | 0 | 0 |
| ML25_014 | 1.59 | 37,774,736 | 75.19 | 28,541 | 12,878 | 48 |
| ML25_015 | 1.77 | 13,085,404 | 38.51 | 3,274 | 1,536 | 7 |
| ML25_016 | 1.94 | 40,345,890 | 77.70 | 15,366 | 6,124 | 28 |
| ML25_022 | 3.03 | 33,927,710 | 73.12 | 13,490 | 3,550 | 16 |
| ML25_026 | 3.87 | 4,246,725 | 62.65 | 958 | 585 | 4 |
| ML34_003 | 0.38 | 18,535,314 | 15.74 | 11,019 | 1,458 | 5 |
| ML34_006 | 0.84 | 19,429,820 | 73.10 | 16,192 | 5,748 | 26 |
| ML34_007 | 0.99 | 20,155,676 | 80.16 | 13,528 | 5,123 | 25 |
| ML34_010 | 1.44 | 20,207,295 | 3.85 | 3,718 | 1,187 | 3 |
| ML34_012 | 1.77 | 28,980,246 | 94.50 | 3,862 | 2,362 | 15 |
| ML34_014 | 2.10 | 19,268,023 | 79.96 | 13,333 | 3,868 | 22 |
| ML34_015 | 2.27 | 26,572,506 | 80.72 | 17,751 | 7,788 | 45 |
| ML34_016 | 2.43 | 44,069,643 | 75.03 | 31,683 | 12,236 | 54 |
| ML34_017 | 2.59 | 39,831,849 | 83.75 | 13,007 | 5,360 | 22 |
| ML34_020 | 3.13 | 21,090,603 | 62.35 | 26,331 | 9,575 | 36 |
| ML34_022 | 3.76 | 16,805,500 | 70.14 | 15,098 | 4,554 | 21 |
| NR18_003 | 0.25 | 17,734,958 | 10.25 | 12,762 | 872 | 3 |
| NR18_005 | 0.45 | 23,358,319 | 73.17 | 18,185 | 6,182 | 38 |
| NR18_007 | 0.65 | 18,924,563 | 72.48 | 16,652 | 6,220 | 29 |
| NR18_010 | 0.95 | 18,483,671 | 70.87 | 15,895 | 7,177 | 33 |
| NR18_011 | 1.05 | 16,063,433 | 73.97 | 14,261 | 4,992 | 28 |
| NR18_012 | 1.15 | 18,525,675 | 84.53 | 12,613 | 5,359 | 28 |
| NR18_015 | 1.51 | 18,780,223 | 4.32 | 5,533 | 214 | 1 |
| NR18_017 | 1.86 | 21,590,172 | 84.44 | 2,485 | 1,306 | 7 |
| NR18_023 | 2.92 | 35,560,955 | 14.00 | 15,853 | 2,880 | 12 |
| NR18_025 | 3.27 | 45,504,942 | 85.73 | 2,969 | 1,443 | 6 |
| NR18_028 | 3.76 | 12,877,162 | 16.17 | 1,692 | 410 | 2 |
| NR3_003 | 0.34 | 5,781,590 | 51.47 | 740 | 497 | 4 |
| NR3_006 | 0.73 | 40,278,890 | 95.24 | 2,983 | 1,606 | 7 |
| NR3_009 | 1.11 | 7,729,451 | 67.07 | 582 | 361 | 3 |
| NR3_012 | 1.57 | 8,307,886 | 33.98 | 458 | 458 | 2 |
| NR3_017 | 2.72 | 2,971,431 | 10.36 | 794 | 16 | 1 |
| NR3_023 | 3.11 | 20,759,186 | 33.60 | 12,394 | 4,172 | 21 |
| NR3_025 | 3.37 | 45,504,942 | 11.75 | 1,388 | 751 | 2 |
| NR3_028 | 3.83 | 12,877,162 | 17.65 | 11,726 | 1,735 | 8 |
| **Total** | **N/A** | **927,847,495** | **N/A** | **441,724** | **160,695** | **757** |

Positive identification of proteins from LC-MS/MS was only possible for 27 out of 43 submitted protein samples. A total of 29,069 proteins were identified (ranging from 1–8,192 per sample, average of 1,077) (**Table 2**), the majority of which were unique (24,033). Only 10,666 proteins (37%) mapped to binned scaffolds. A major issue impacting data collection was the presence of polyethylene glycol (PEG), a common polymer contaminant in mass spectrometry-based analyses (Keller et al., 2008), which presents as numerous peaks with masses increasing sequentially by 44 kDa, swamping true biological signals (Ahmadi and Winter, 2018). Avoiding PEG contamination is challenging, as it is ubiquitous in many commercial and laboratory products, such as cosmetics, detergents and plastics (Ahmadi & Winter, 2018). Attempts to limit contamination during protein extraction by using HPLC-grade solvents, rinsing glassware with isopropanol instead of soap, and avoiding the use of PEG-containing plastics led to marginal improvements in some samples, but significant PEG contamination was still detected in others. It is possible that PEG was introduced to samples from handling prior to protein extraction, such as during tailings core processing. Additional processing with detergent removal and desalting columns were performed at the BioCORE facility (University of Western Ontario, Canada, see Methods), but these steps did not result in an improvement on the observed relative peptide signal.

**Table 2:** Metaproteomic sequencing statistics by sample. Peptides were identified using LC-MS/MS and mapped to a non-redundant database of predicted proteins from metagenomic sequencing data. The total number of potentially expressed proteins within a sample is reported as the Protein # along with the total number of Peptides supporting the identified proteins (Peptide #). A minimum of one unique peptide within a protein group (accounting for homologs and variable isoforms) and a false-discovery rate of 1% was used as the threshold for positive protein identification.

| Sample | Depth (m) | Protein # | Peptide # | Unique Peptides |
| --- | --- | --- | --- | --- |
| ML25_003 | 0.28 | 8 | 8 | 4 |
| ML25_006 | 0.62 | 8,192 | 20,044 | 6,796 |
| ML25_007 | 0.73 | 2,065 | 2,866 | 801 |
| ML25_009 | 0.96 | 737 | 759 | 660 |
| ML25_010 | 1.07 | 532 | 537 | 445 |
| ML25_013 | 1.42 | 6 | 6 | 4 |
| ML34_003 | 0.38 | 179 | 193 | 66 |
| ML34_006 | 0.84 | 291 | 403 | 84 |
| ML34_007 | 0.99 | 2,036 | 3,833 | 1,888 |
| ML34_010 | 1.44 | 2,770 | 3,964 | 1,024 |
| ML34_012 | 1.77 | 5,470 | 11,675 | 3,863 |
| ML34_014 | 2.1 | 9 | 9 | 7 |
| ML34_015 | 2.27 | 61 | 68 | 17 |
| ML34_016 | 2.43 | 20 | 22 | 7 |
| ML34_017 | 2.59 | 1 | 2 | 2 |
| ML34_020 | 3.13 | 365 | 550 | 252 |
| ML34_022 | 3.76 | 47 | 52 | 12 |
| NR18_003 | 0.25 | 58 | 62 | 32 |
| NR18_010 | 0.95 | 204 | 286 | 69 |
| NR18_011 | 1.05 | 734 | 979 | 250 |
| NR18_012 | 1.15 | 65 | 70 | 7 |
| NR18_015 | 1.51 | 206 | 208 | 46 |
| NR18_023 | 2.92 | 585 | 591 | 464 |
| NR18_025 | 3.27 | 2 | 2 | 2 |
| NR3_009 | 1.11 | 489 | 494 | 418 |
| NR3_023 | 3.11 | 548 | 558 | 491 |
| NR3_028 | 3.83 | 3,389 | 5,069 | 2,342 |
| **Total** | **N/A** | **29,069** | **53,309** | **20,053** |

The low number of MAGs (compared to ASVs from the same samples) and identified proteins in this study reflect the ongoing challenges in obtaining high-quality DNA and protein samples from mine tailings. At baseline, mine tailings exhibit many of the same issues as soil for DNA and protein extractions, including the presence of high molecular weight humic acids and recalcitrant clays (Pan *et al*., 2024). Further, low biomass and contaminants within the sample matrix (*e.g.*, heavy metals) limit the effectiveness of current DNA/protein extraction methods, which are optimized for soil communities. Loq biomass protein samples will also be sensitive to contaminants introduced during sample processing (*e.g.*, PEG), resulting in low signal-to-noise ratios in mass spectrometry. Nevertheless, these paired metagenome/metaproteome samples allowed for examination of the abundant members of the tailings microbial communities, and, for some samples, a robust determination of highly expressed genes via proteomics. Though less comprehensive than initially hoped, this still represents a unique dataset of DNA and protein extracted directly from mine tailings.

### Microbial community overview

Community profiling based on metagenomic sequencing data was low resolution due to the limited number of high-quality MAGs. In some cases, a sample appeared to be comprised of only 1–2 populations (**Figure 1**), but it was clear from 16S rRNA amplicon sequencing (Chen et al., 2024) that not all of the community diversity was captured in the assembled and binned scaffolds, especially for lower-abundance organisms. Consequently, quantitative comparisons of relative MAG abundances in this study (**Figure 1**) should be viewed with caution, especially for samples with low bin counts (<10). However, the presence and localization of abundant taxa (>15%) generally agreed between the two datasets (**Figure 1**) (Chen et al., 2024).

**Figure 1:**
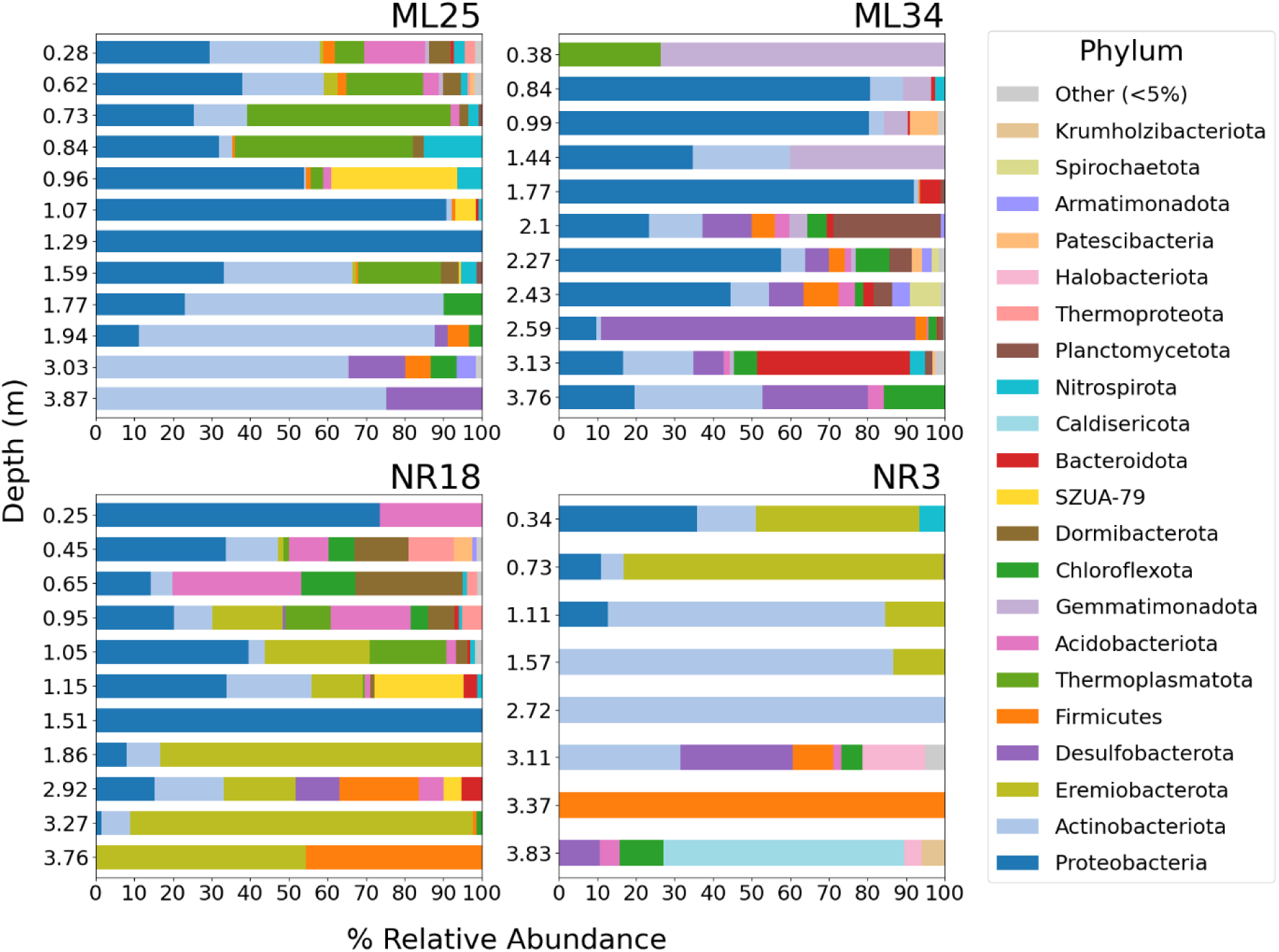
Microbial community composition across tailings cores from metagenomic sequencing data. Bars indicate phylum-level relative abundances of MAGs within each sample, based on the average depth of coverage of scaffolds within individual MAGs. Only phyla present at a relative abundance >5% in at least one sample in any dataset are shown. Phyla are ordered from most overall abundant (left) to least abundant (right), or bottom-to-top in the legend. MAGs were classified with GTDB-tk (v2.1.0, database release version 207.0). Note that nomenclature for the Proteobacteria (Pseudomonadota) and Firmicutes (Bacillota) have been updated in more recent versions of the GTDB. The y axes indicate the sample depth below ground level and are not scaled proportionally.

In terms of phylogenetic diversity, the Proteobacteria (mainly Gammaproteobacteria) and Actinobacteriota phyla had the highest overall abundance (**Figure 1**) and the highest number of non-redundant MAGs across samples (**Figure 2**). Most of the identified proteobacterial MAGs belong to clades associated with iron and/or sulfur oxidation in mining waste, but there were differences in the localization of individual taxa. Within the Acidiferrobacterales, *Acidiferrobacter* spp. were unique to ML25 whereas *Sulfuricaulis* spp. were almost exclusively found in NR18. *Sulfuricella*, *Sulfuriferula*, and *Thiobacillus* spp. (o. Burkholderiales) were mainly found in ML25 and ML34, with *Thiobacillus* much more abundant in ML34 samples. *Acidithiobacillus* spp. were found in all four tailings locations, but *A. ferrooxidans* were highly abundant in ML25, whereas *A. ferrivorans* was more common in other samples.

**Figure 2:**
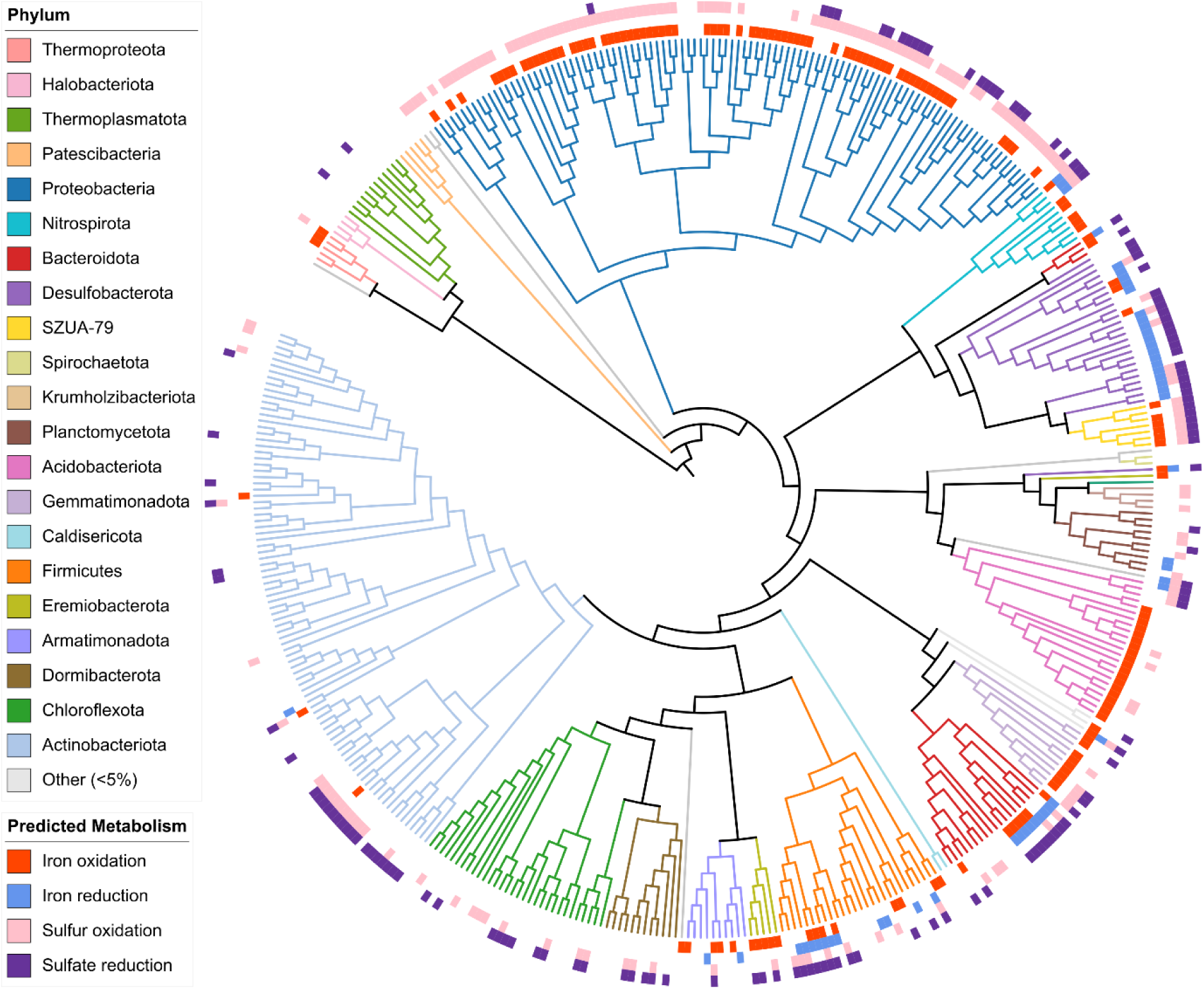
Phylogenetic distribution of sulfur and iron cycling genes across all MAGs. Tree branches are unscaled (cladogram) and colored by GTDB-tk assigned phylum in clockwise order following the legend, starting from the archaea (top left). Only phyla present at a relative abundance >5% in at least one sample in any dataset are colored. Outer rings represent presence/absence of iron and sulfur cycling genes, where at least one hit for any gene within each category was denoted as presence. Iron and sulfur metabolism genes were compiled from FeGenie, DRAM, and DiSCo annotations. Iron oxidation genes include *cyc1*/*cyc2*, *fox* genes, and sulfocyanin; reduction genes include *omc* and *dfe* genes. Sulfur oxidation is represented by genes in the *sox* or oxidative type *dsr* pathways; reduction genes include thiosulfate reductase, tetrathionate reductase, and reductive type *dsr* genes.

Unlike the Proteobacteria, Actinobacteriota MAGs were relatively novel, with most unclassified at the family or genus level, and with only one mining waste-associated Actinobacteriota (*Ferrimicrobium* spp.) identified, and only at <4% abundance. The majority of Actinobacteriota in shallow samples (<2 m) belong to the Acidimicrobiales order, and appear to be mostly organoheterotrophs. Deeper samples (>2 m) included members of the understudied anaerobic Thermoleophilia and Humimicrobiia clades, some of which are predicted to be sulfur oxidizers and/or sulfate reducers based on *sox* and/or *dsr* gene annotations (**Figure 2**).

Other abundant lineages mapping to predicted roles in iron and/or sulfur cycling included the Desulfobacterota, Acidobacteriota (ML25/NR18), Bacteroidota (ML34), Firmicutes (NR18/NR3), SZUA-79 (ML25/NR18), and Eremiobacterota (NR18/NR3), which are discussed in that context below. Abundant phyla that did not contribute to the observed iron/sulfur cycling gene abundances or gene expression included the archaea (Thermoplasmatota), Caldisericota, Planctomycetota, Gemmatimonadota, and Dormibacterota.

### Distribution and expression of MRGs

Annotating MRGs using the built in BacMet-scan software associated with the BacMet bacterial metal and biocide resistance gene database (Pal et al., 2014) returned 329,520 non-redundant hits from all binned scaffolds. Annotation using custom Hidden Markov Models (HMMs) built from the same BacMet database (see Methods) returned 1,660,455 hits at the same level of statistical significance (E<0.05) (**Table S1**). The BLAST-based approach used by BacMet-scan is generally considered less sensitive than HMMs (Krogh et al., 1994; Park et al., 1998), where HMMs can identify more evolutionarily distant homologs across gene families, enabling the detection of a broader range of genes (Park et al., 1998).

Given these advantages, we proceeded with data analysis using results from the BacMet HMM-based gene annotations. The HMM set excluded genes with <5 representative sequences in the BacMet database (84 out of 613 unique genes). To partially account for this, the BacMet-scan BLAST hits for these 84 genes were appended to the HMM annotations, which added 10,462 additional hits, or a 0.6% increase. Both methods lack the ability to annotate archaeal genomes as the BacMet database only includes bacterial sequences. This means the 22 archaeal MAGs of the 454 total non-redundant MAGs (4.8%) were not included in this analysis.

The overall abundance of MRGs was generally consistent across samples within each tailings core, with two exceptions in NR18 (1.51 m) and NR3 (3.37 m), both showing elevated proportions across MRGs (**Figure 3**). These samples had poor assembly and binning statistics, with only 3 total high-quality MAGs, (**Table 1**) and thus MRGs encoded on these MAGs are likely over-represented compared to the complete microbial communities. When comparing gene abundances by metal, zinc MRGs were the most common, followed by copper, nickel, and iron (**Figure 3**). Most MRGs associated with zinc exhibit broad specificity and also confer resistance to other metals (most commonly cadmium and copper) (Pal et al., 2014), which could explain the overrepresentation of zinc MRGs as these MRG hits would be counted multiple times, despite zinc not being a major contaminant in these tailings (<9.0 mg/L) compared to the levels of iron, nickel, and copper at these sites (**Supplemental Data File 1: Table S2**).

**Figure 3:**
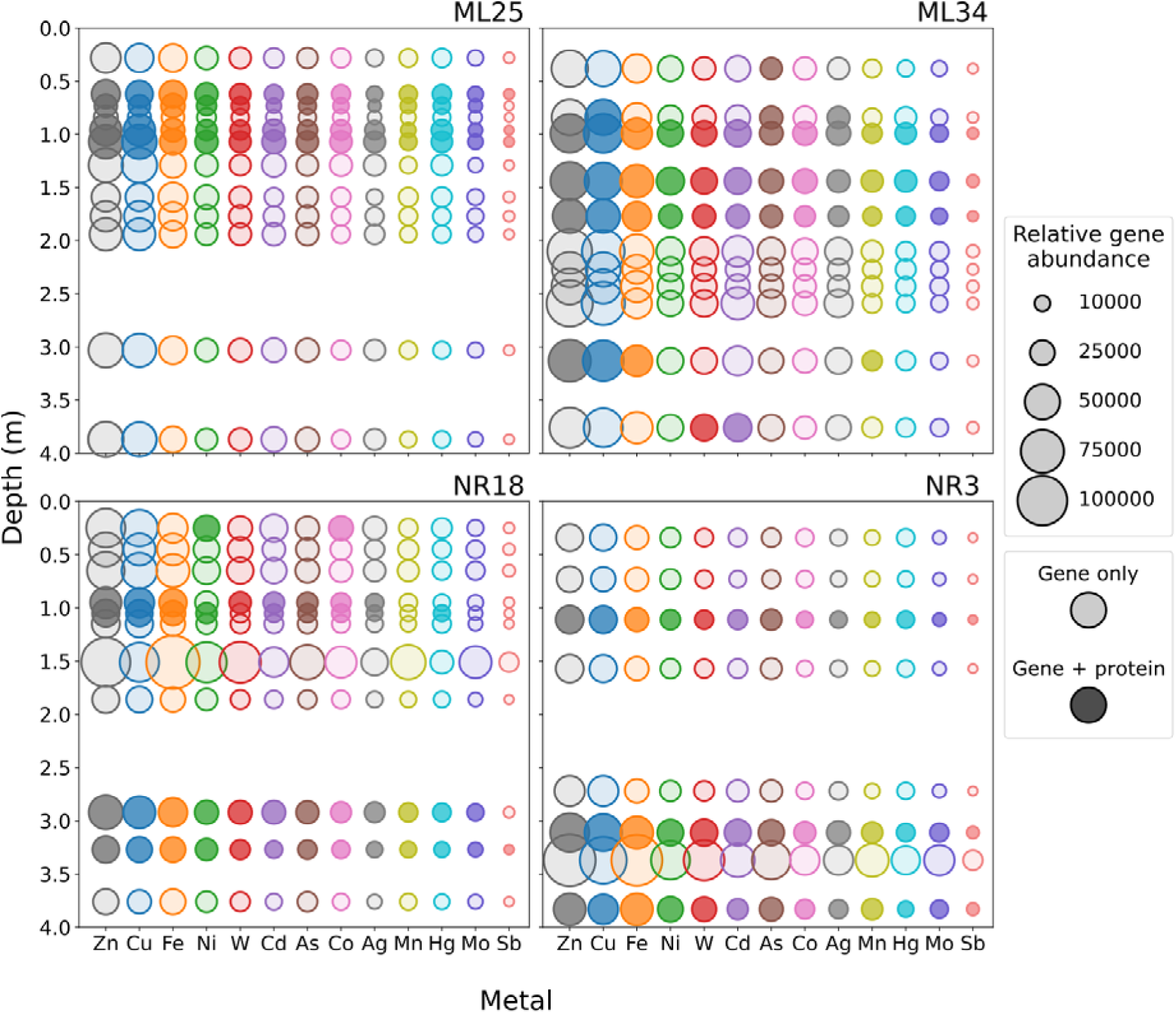
Distribution of MRGs across vertical mine tailings transects. MAGs were annotated using profile HMMs of genes in the BacMet database. Rare genes (<5 representatives in the BacMet database) were annotated with BacMet-scan. Circle diameters representing relative gene abundances were calculated as the number of hits multiplied by the percent relative abundance of the MAG associated with each hit. In samples where protein expression was detected, the corresponding gene(s) are indicated by circles with darker shading. MRGs are colored based on metal, ordered from highest overall abundance (Zn) on the left to lowest (Sb) on the right. Multi-metal resistance genes were counted towards multiple categories. Only categories with abundances >2% of the total are shown.

Differences in metal toxicity may also affect the relative abundance of their associated MRGs. Resistance mechanisms for highly toxic metals such as arsenic and mercury are likely to be encoded in microbial genomes regardless of whether environmental concentrations are high, as these pathways evolved early in Earth’s history and are widely distributed among bacteria and archaea (Osborn et al., 1997; Zhu et al., 2017). Likewise, copper resistance genes were more abundant than nickel and iron resistance genes (**Figure 3**), even though copper concentrations were lower in most samples, likely reflecting copper’s greater cytotoxicity relative to iron and nickel (Fait et al., 2006; Gilani & and Alibhai, 1990).

There was no significant correlation between individual concentrations of major heavy metals (Cu, Ni, Fe) and their associated MRG abundance (Pearson correlation, p_Cu_=0.18, p_Fe_=0.79, p_Ni_=0.28). More heavily contaminated tailings (*e.g.*, ML25) had similar or lower Cu, Ni, and Fe resistance gene abundances to other locations (**Figure 4, Figures S1-S3**). There was also no correlation between metal concentrations and MRG abundances when comparing samples within a core (0.05≤p≤0.95), a slight positive correlation between total MRG abundances with pH (r=0.54, p=0.003), and a slight negative correlation between total MRG abundances with redox potential (r=-0.48, p=0.009), contradictory to the expectation that metals are more bioavailable and mobile at lower pH and higher redox potential, therefore leading to higher MRG abundances. Although higher environmental metal concentrations could select for organisms with more resistance genes targeting those metals, the results from this study suggest that a broad range of MRGs are present in mine tailings microbial communities regardless of the level of heavy metal stress they are currently experiencing.

**Figure 4:**
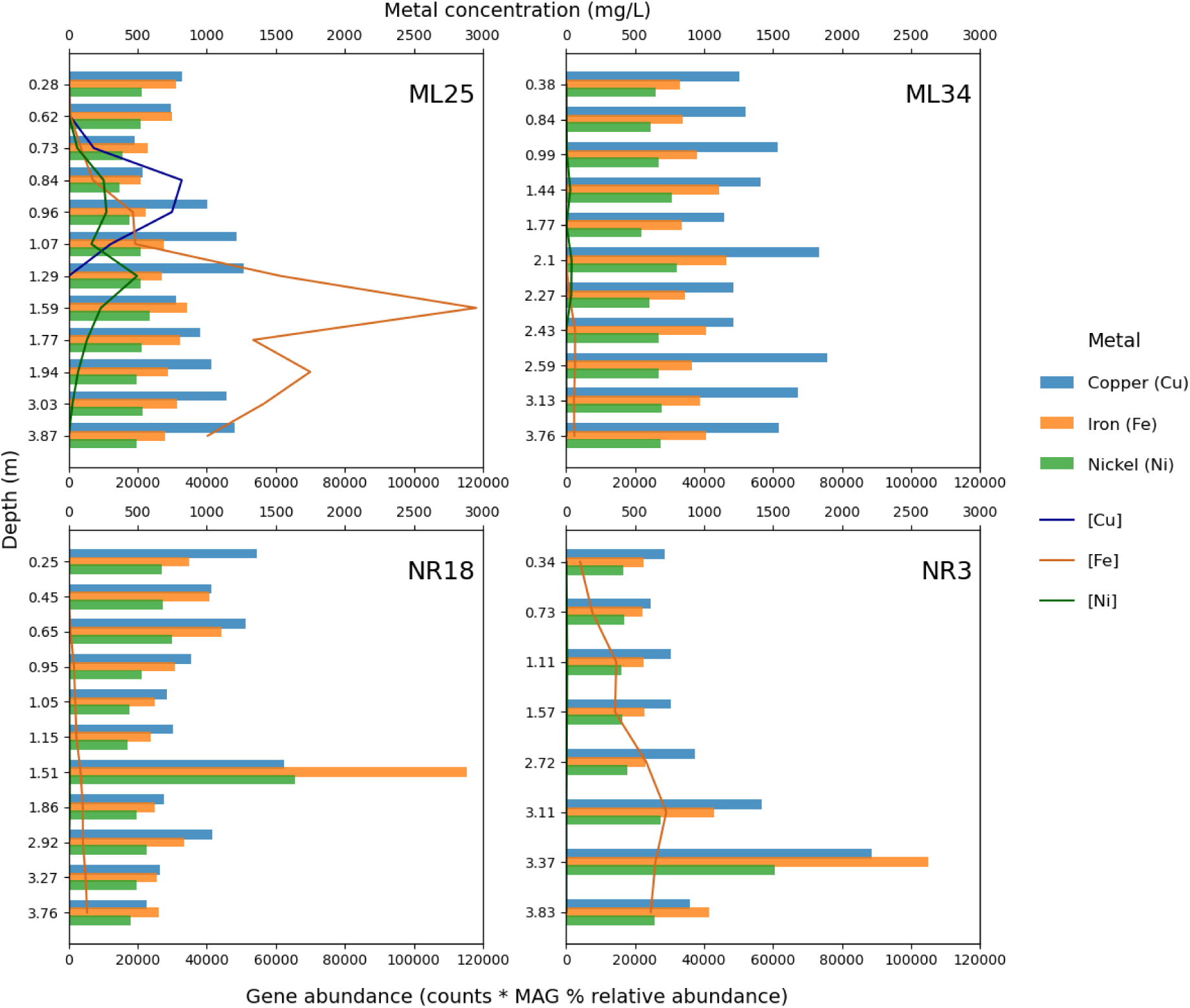
Abundance of metal resistance genes and metal concentrations along vertical mine tailings transects. Bar plots depict relative gene abundances for Cu, Fe, and Ni-associated MRGs. Line plots represent Cu, Fe, and Ni concentrations in tailings pore-water. There was no significant correlation between metal concentrations and associated MRG abundances (p_Cu_=0.18, p_Fe_=0.79, p_Ni_=0.26). Note that the y axes are not scaled proportionally to depth.

Metal resistance in mine tailings communities could instead be regulated at the gene expression level (Chandrangsu et al., 2017; Nies & Brown, 1998), which cannot be directly inferred from metagenomic sequencing data. In samples with high (>2000) protein counts, resistance genes for most metals were expressed (**Figure 3**), confirming that many MRGs are not only encoded in genomes, but also actively expressed by microbes inhabiting tailings environments. Absence in other samples, especially those with few identified proteins, cannot be confidently interpreted as a lack of MRG expression (**Figure 3**). Correlations between MRG expression levels and extracellular metal concentrations could not be determined due to the lack of quantitative mass spectrometry data. However, we hypothesize that the same factors influencing MRG abundance in metagenomes, such as the relative toxicity of individual metals and the ability of MRGs to confer multi-metal resistance, are important determinants of protein abundance.

Some MRGs are known to be carried on plasmids (Collard et al., 1994; Foster, 1983; Li et al., 2015) as opposed to genomic DNA, leading to their potential underrepresentation when only considering MRGs associated with MAGs. We performed an additional analysis to investigate the contribution of MRGs associated with unbinned scaffolds, including mobile genetic elements, relative to the total MRG abundance within the community. We ran the BacMet HMM annotation pipeline again, on all scaffolds >2.5 kbp in length, and hits were mapped to four categories based on the scaffold of origin: binned, unbinned chromosomal, unbinned plasmids, and unbinned phages as defined by PPRmeta (Fang et al., 2019) (see Methods). When considering the total MRG counts across scaffolds, 40.1% (934,294 out of 2,328,173) were located on unbinned scaffolds, with 14.6% of the total associated with plasmids (**Figure S4, Supplemental Data File 1: Table S3**). However, when normalizing MRG hits to relative scaffold abundances, only 19.4% of MRGs are associated with unbinned scaffolds (5.9% plasmid) (**Figure S4**, **Supplemental Data File 1: Table S4**). These findings support that the majority of MRGs present in the community that could be sequenced and assembled were captured in the binned scaffolds, and that plasmid-associated MRGs likely play a relatively minor role in conferring metal resistance in tailings.

To investigate the metal resistance mechanisms of individual bins, MAGs were ranked by the total number of predicted Cu, Ni, and Fe resistance genes they encoded (**Figure S5**), as high MRG counts may be indicative of high metal tolerance in an organism. MAGs encoding a high number of copper-resistance genes (top 200) were most commonly members of the Proteobacteria (30.5%), Desulfobacterota (12.5%), Chloroflexota (10.0%), and Acidobacteria (7.5%) (**Figure S5**). The top 5 MAGs with the highest number of predicted copper-resistance genes were taxonomically novel (**Table 3**), unclassified at the genus level or above, and may represent interesting targets for future isolation and characterization of copper resistance mechanisms.

**Table 3:** Metagenome-assembled genomes from mine tailings with the top five highest (99^th^ percentile) copper, nickel, and iron resistance gene counts. MAGs (Bin ID) are ranked in decreasing order of MRG counts (number of genes). The lowest assigned level of GTDB taxonomic classification is reported (c=class, o=order, f=family). MAG relative abundances within their sample (Rel. Abundance) were calculated based on average scaffold depth of coverage.

| Metal | Bin ID | Taxonomic classification | # of genes | Rel. Abundance (%) |
| --- | --- | --- | --- | --- |
| Copper | ML34_020_30 | f. Myxococcaceae | 1,590 | 0.96 |
|  | ML34_015_36 | c. Anaerolineae | 1,363 | 0.68 |
|  | NR18_007_01 | o. Aggregatilineales | 1,250 | 1.80 |
|  | ML34_022_13 | o. Deferrisomatales | 1,222 | 1.02 |
|  | NR3_023_07 | f. Pseudopelobacteraceae | 1,123 | 7.38 |
| Nickel | ML34_015_29 | *Allorhizobium* sp. | 701 | 0.94 |
|  | NR18_015_01 | *Bradyrhizobium* sp. | 646 | 100 |
|  | NR3_025_02 | *Ruminiclostridium* sp. | 622 | 28.87 |
|  | NR3_023_12 | *Mobilitalea* sp. | 608 | 1.85 |
|  | NR18_005_37 | o. Acidimicrobiales | 594 | 1.96 |
| Iron | NR3_025_02 | *Ruminiclostridium* sp. | 1,197 | 28.88 |
|  | ML34_015_29 | *Allorhizobium* sp. | 1,164 | 0.94 |
|  | NR18_015_01 | *Bradyrhizobium* sp. | 1,153 | 100 |
|  | NR3_023_12 | *Mobilitalea* sp. | 1,117 | 1.85 |
|  | NR18_005_37 | o. Acidimicrobiales | 1,103 | 1.96 |

Compared to Cu resistance, there was more taxonomic overlap between MAGs with high nickel-and high iron-resistance gene counts (**Figure S5**), as many genes are mapped to both functions in the BacMet database (Pal et al., 2014). Most of the top 200 highest nickel- and iron- resistance gene-encoding MAGs were classified to members of the Actinobacteriota (28.3%), Proteobacteria (14.75%), Chloroflexota (13.75%), and Firmicutes (9.23%) (**Figure S5**). The top 5 MAGs with highest counts for nickel and for iron-associated MRGs were identical, and included *Allorhizobium* and *Bradyrhizobium* spp. (c. Alphaproteobacteria, o. Rhizobiales) (**Table 3**). *Bradyrhizobium* and other Rhizobiales spp. are known to be nickel tolerant and promote plant growth in Ni-rich soils, as they are nitrogen-fixing symbionts of legumes (Chaintreuil et al., 2007; Liu et al., 2022). These lineages potentially enhance tailings remediation through Ni uptake and bioaccumulation, thereby decreasing its mobility and impact on other community members (Maier et al., 1990; Stults et al., 1987). The *Ruminiclostridium* MAG in NR3_025 is artificially abundant, as only 2 MAGs were binned for that sample. *Ruminiclostridium* may be present due to the presence of organic plant material in deep NR3 samples, as it is expected to use an anaerobic, cellulose degrading metabolism (Rettenmaier et al., 2021).

The MAGs high in Cu, Ni, and Fe resistance genes are predicted to represent heterotrophic organisms (**Table 3**), giving a diverse pool of potential metal cyclers. However, the majority of detected metal resistance proteins mapped to *Acidithiobacillus*, *Thiobacillus*, and other autotrophic lineages (**Figure S6**), with the exceptions of *Bradyrhizobium, Metallibacterium, Methylovirgula,* and *Devosia* spp., where a much smaller proportion of the predicted metal cyclers were actively expressing these functions. Of these, a *Thiobacillus* spp. from ML34 appeared to express the largest diversity of unique proteins related to copper, nickel, and iron resistance (**Figure S6**). Fewer types of metal resistance proteins were detected from the taxa predominantly inhabiting more highly metal-contaminated tailings such as *Acidithiobacillus*, *Metallibacterium*, and Acidiferrobacteraceae, although specific proteins/complexes such as Cme, Cus, Czn, Nik, and Nia were commonly detected as expressed by these clades. From these data we hypothesize that microbial adaptations to heavy metal stress more frequently involve the upregulation of a few select pathways for metal detoxification as opposed to acquiring and utilizing many different pathways.

### Distribution and expression of iron- and sulfur-related genes

In general, iron- and sulfur- oxidation genes were more abundant in shallow samples whereas reductive metabolisms were more abundant in deeper samples (**Figure 5**, **Figure 6**), which was expected based on decreasing oxygen availability and redox potentials (**Table S2**) with increasing depth. Most predicted iron-oxidizers were bacteria using the *cyc2* pathway (Castelle et al., 2008; McAllister et al., 2020). Iron reduction genes were more diverse, being evenly distributed among hits for *dfe_0461-0465*, *omcF/S/Z*, and other pathways (Deng et al., 2018; Santos et al., 2015). There was considerable overlap between predictions of iron oxidizing and sulfur oxidizing Gammaproteobacteria, as well as between iron reducing and sulfate reducing Desulfobacterota (**Figure 2**). As seen for the MRGs, corresponding proteins were only detected in a few samples (**Figure 5**, **Figure 6**). In many cases, some genes for a pathway were detected as proteins but not others, and this was inconsistent between samples from similar depths in the same tailings core. Due to this low coverage, protein data does not represent a comprehensive profile of microbial gene expression in these tailings, but provides information as to active processes at the time of sampling.

**Figure 5:**
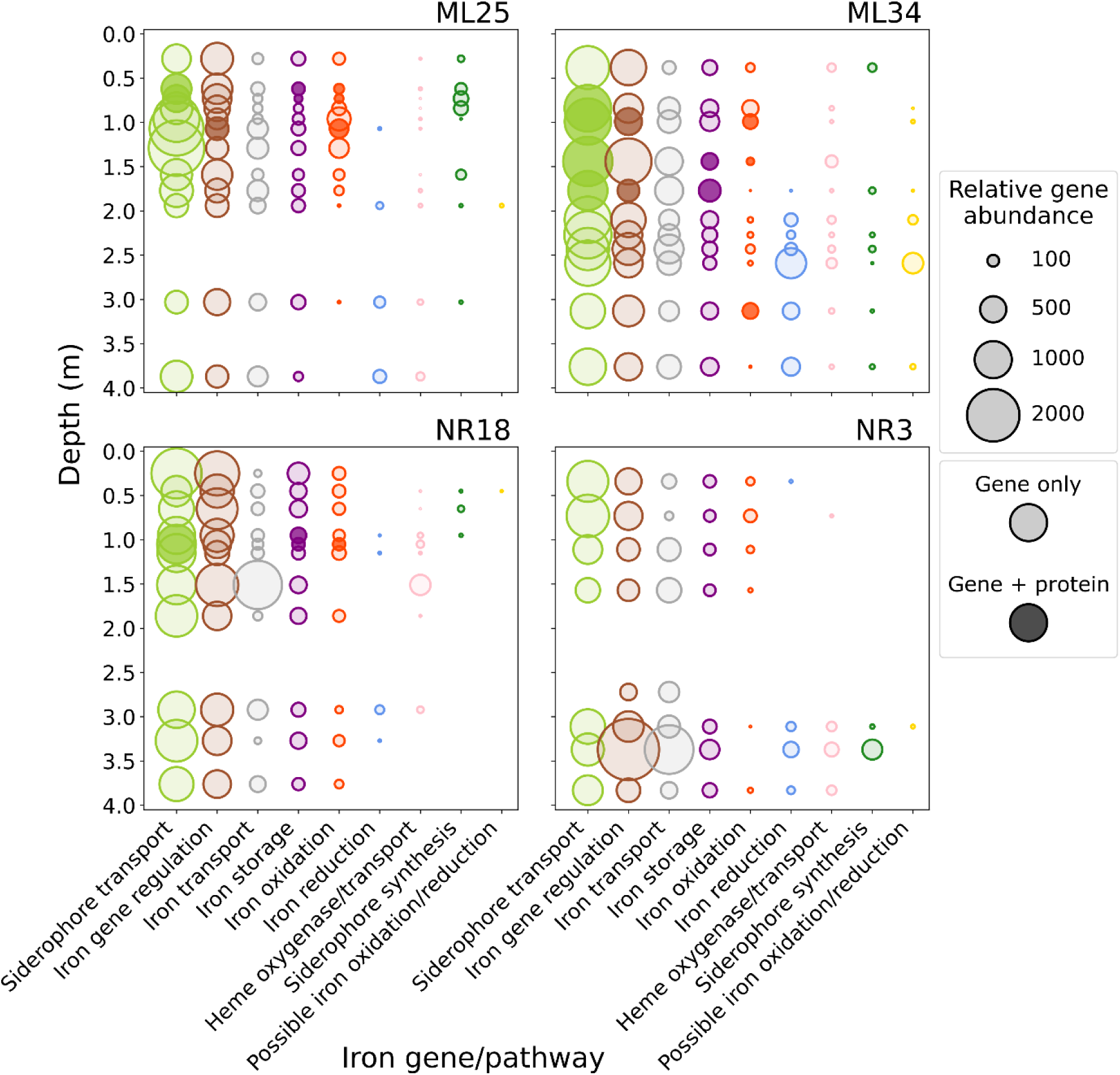
Distribution of iron-related genes across vertical tailings transects. MAGs were annotated using FeGenie. Relative gene abundances were calculated as the number of hits multiplied by the percent relative abundance of MAGs associated with each hit. In samples where protein expression was detected, the corresponding gene(s) are indicated by circles with darker shading. Gene categories are colored based on function (**Table S5**) (Garber et al., 2020), ordered from highest overall abundance (Siderophore transport) on the left to lowest (Possible iron oxidation/reduction) on the right.

**Figure 6:**
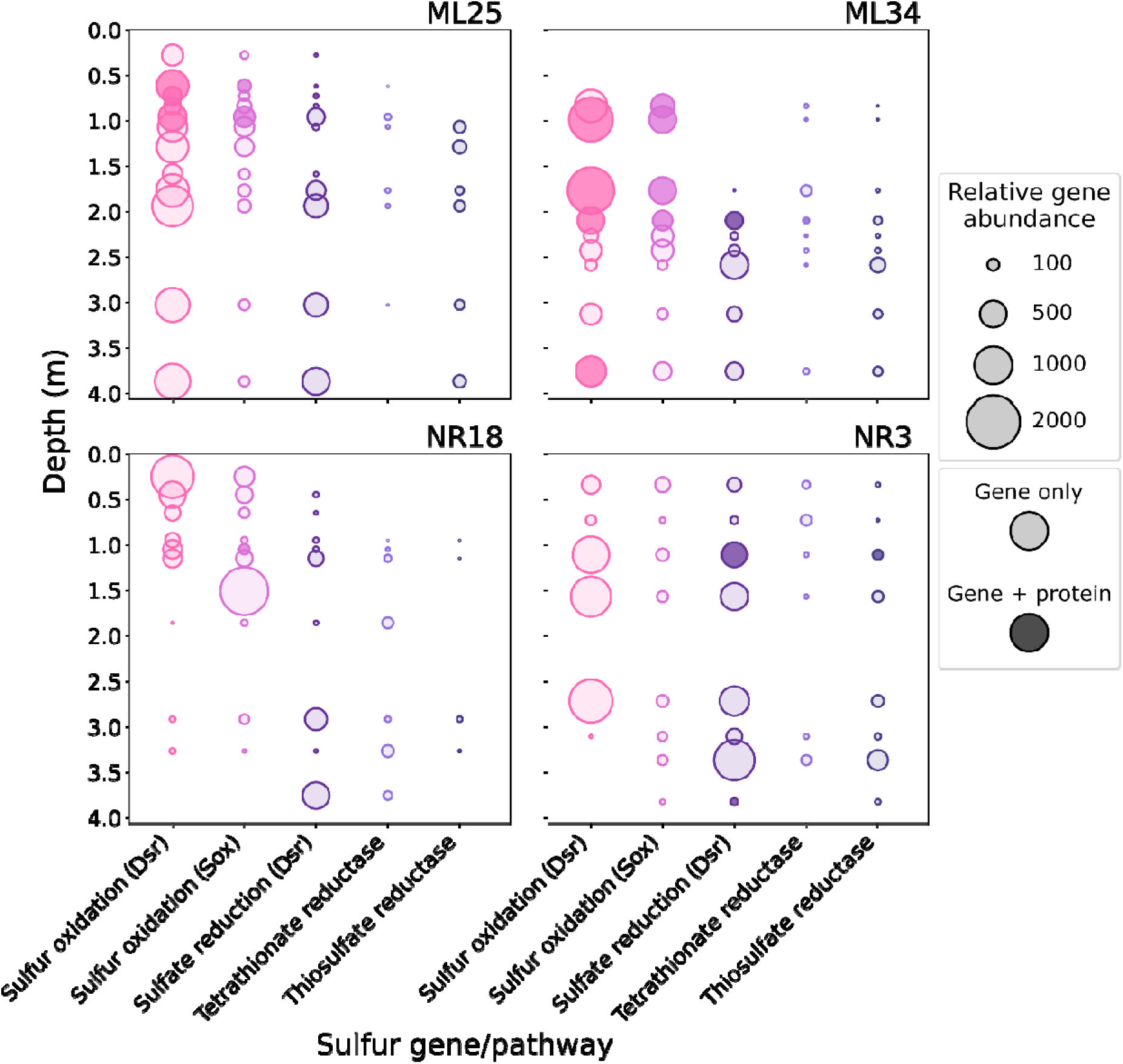
Distribution of sulfur-cycling genes across vertical tailings transects. MAGs were annotated using DRAM (*sox*, tetrathionate reductase, and thiosulfate reductase genes) and DiSCo (*dsr* genes). Relative gene abundances were calculated as the number of hits multiplied by the percent relative abundance of MAGs associated with each hit. In samples where protein expression was detected, the corresponding gene(s) are indicated by circles with darker shading. Gene categories are colored based on function, ordered from highest overall abundance (Sulfur oxidation, Dsr pathway) on the left to lowest (Thiosulfate reductase) on the right.

Differences in potential iron oxidation and *sox*-based sulfur oxidation were most pronounced in ML25, with gene abundances gradually increasing until 1.0–1.5 m, where the microaerophilic environment favored biotic pyrite oxidation, then gradually decreasing as the environment shifted anoxic (**Figure 5**, **Figure 6**). This trend was mostly attributed to *Acidithiobacillus ferrooxidans* (Fe & S), *Acidiferrobacter* (Fe & S), *Sulfuriferula* (S), and *Sulfuricella* spp. (S), although many other predicted Fe and/or S oxidizers were also present, including *Leptospirillum* (Nitrospirota; Fe), *Metallibacterium* (Proteobacteria; Fe), *Thiomonas* (Proteobacteria; S), unclassified Acidobacteriota MAGs (mainly Fe, some Fe & S), Firmicutes MAGs (varied), *Ca.* Acididesulfobacter (SZUA-79; Fe & S), and *Ca.* Acidulodesulfobacterium spp. (SZUA-79; Fe & S).

The *Ca.* Acidulodesulfobacterales (SZUA-79) MAGs were especially abundant around 1.0 m in depth in ML25 and NR18 (up to 32.6% and 23.0%, respectively). Members of this lineage have been characterized from metagenomic sequencing data from one other AMD environment, with similar hypothesized roles in Fe/S cycling (Tan et al., 2019), but there is currently no isolated representative for this order. Cyc2 (iron oxidation) protein expression from both *Ca*. Acidulodesulfobacter and *Ca*.

Acidulodesulfobacterium was detected (**Figure S7**), and these Cyc2 proteins were most similar to the Leptospirillaceae Cyc2 variant based on FeGenie HMMs. Dissimilatory sulfate reduction activity was also detected from several SZUA-79 MAGs (**Figure S8**), overlapping imperfectly with Cyc2-expressing MAGs (**Figure S7**). Iron oxidation and sulfate reduction are not compatible in the same redox environment; proteins from both pathways were detected from each of the SZUA-79 MAGs in the ML25_007 sample (0.73 m), but it is unlikely that sulfate reduction is supported at this depth. However, these results constitute evidence of *Ca*. Acididesulfobacter and *Ca*. Acidulodesulfobacterium being capable of both metabolisms depending on the surrounding geochemical conditions and the availability of different electron acceptors.

Siderophore-related gene abundance followed similar depth patterns as iron oxidation gene abundance in ML25 (**Figure 5**), as 95% of predicted iron oxidizers co-encoded siderophore genes. Some siderophore transport proteins were detected, but not proteins for siderophore synthesis, in keeping with geochemical conditions: soluble iron is unlikely to be limiting in tailings environments.

Lower acidity in ML34 tailings supported populations of neutrophilic Fe and/or S oxidizers such as *Thiobacillus*, *Halothiobacillus*, and *Gallionella* spp. Proteomic data indicate that *Thiobacillus* were using both Dsr and Sox pathways for sulfur oxidation (**Figure S8**), and that this genus accounted for the majority of iron storage proteins (ferritin) detected (**Figure S7**). Given that *Thiobacillus* cannot oxidize iron for energy, they most likely use ferritin as an iron resistance mechanism (Smith, 2004) by storing excess iron released from oxidizing pyrite and other minerals containing both iron and reduced inorganic sulfur compounds (RISCs). The expression of FecR, a protein regulating the ferric citrate uptake system, was associated with *Ca*. Devosia spp. in ML34 (**Figure S7**), which also expressed a variety of Cu, Ni, and Fe-resistance proteins (**Figure S6**). Members of this genus are often found in soil environments contaminated with various organic toxins (Talwar et al., 2020); this study demonstrates their tolerance towards moderate levels of metal contamination as well.

The *Ca.* Eremiobacterota were predicted iron oxidizers unique to NR18 and NR3 samples, and were the most abundant organisms (up to 89%) in several samples. All *Ca.* Eremiobacterota MAGs were classified under Ca. Baltobacteraceae, a family whose members are predicted to be metabolically versatile based on previous environmental sequencing data (Ji et al., 2021). Members of the Baltobacteraceae have not previously been identified in mining-associated environments, nor have they previously been predicted to capable of autotrophic iron oxidation (Ji et al., 2021; Pessi et al., 2024; Ward et al., 2019; Yabe et al., 2022).

The majority (68%) of predicted iron-reducing organisms (mostly from the Desulfobacterota) were identified from ML34 samples below 1.5 m in depth (**Figure 5**). None of the predicted iron reducers were classified below the family level. No iron reducing gene expression was detected in the proteomics, although this may be due to the fact very few proteins were identified from deeper samples, especially from the ML34 core (**Table 2**). An abundant (37%) Ignavibacteriaceae (Bacteroidota) MAG in the ML34_020 (3.13 m) sample encoded both iron oxidation and reduction genes, metabolisms which have not been ascribed to the only isolated representative in this genus (Iino, 2014). Cyc2 expression was detected in this sample (**Figure 5**), but the proteins mapped to *Gallionella*, *Ca.* Acidulodesulfobacterium, and unbinned scaffolds rather than to this Ignavibacteriaceae MAG. The geochemical conditions at 3.13 m depth are unlikely to be supporting significant rates of iron oxidation.

Predicted sulfate-reducing organisms were generally localized to deeper (> 2 m) samples at all locations, with some exceptions (such as *dsr* encoded and/or expressed by *Thiobacillus*) (**Figure 6, Figure S8**) possibly due to misclassification of oxidative *dsr* genes as reductive. Deeper samples generally had poor binning and protein coverage, and a comprehensive profile of all sulfate reducers cannot be described from these samples. Nonetheless, many novel taxa that were unclassified at the genus level and above were found to both encode and express genes for dissimilatory sulfate reduction. These include members of the Firmicutes (Desulforudaceae, Desulfotomaculales), Desulfobacterota (Desulfobulbaceae, Desulfobaccales, Desulfurivibrionaceae, Desulfomonilaceae), SZUA-79 (discussed above), and one Actinobacteria MAG (Solirubrobacteraceae) (**Figure S8**). Biological sulfate reduction is often targeted in remediation of heavy metals from mining waste, as heavy metals can be immobilized via sulfide precipitation (Johnson & Hallberg, 2005). Although this method is typically not applicable for remediation of tailings at shallow depths, the presence of sulfate reducing organisms at these depths suggested that the remedial organic carbon cover is effectively removing oxygen, thereby limiting further oxidation of deeper sulfide minerals. The diverse set of novel predicted sulfate reducers identified in these tailings warrant further investigation into their ability to control metal mobility *in situ*, as well as whether measures can be taken to stimulate their activity, such as the addition of organic carbon, nitrogen, and/or phosphorus sources. Improvements in DNA and protein extraction are also required to provide more comprehensive multi-omic analyses of novel sulfate reducers and their capacities for metal resistance.

A wide variety of taxa appear to be capable of Fe/S cycling in tailings. Vertical heterogeneity in tailings support niches for Fe/S oxidation and reduction, but the exact populations occupying these niches can be highly variable between different cores, and taxonomic composition of these guilds is likely based on selection from environmental factors such as pH and contaminant profiles (Chen et al., 2024).

Metagenomics enabled the prediction of 301 potential Fe/S cycling MAGs (**Figure 2**), of which 139 (46.2%) corresponded to ASVs that could not be confidently linked to these metabolisms based on taxonomic classifications alone (Chen et al., 2024). Although most iron- and sulfur-oxidizers identified in this study belong to well-studied genera with cultured representatives (**Figure S7, Figure S8**), there was considerable species and strain level functional diversity within these groups that potentially influence their roles in heavy metal cycling and/or applicability towards bioleaching (Issotta et al., 2018; Kumar et al., 2019; Orell et al., 2010; Zhang et al., 2016). Furthermore, organisms encoding and/or expressing proteins for dissimilatory sulfate reduction were largely unclassified at the genus level or above (**Figure S8**), and were missed in functional predictions from 16S rRNA data (Chen et al., 2024). It is clear from this study that a considerable amount of yet uncharacterized diversity remains within Fe/S cycling consortia found in mine tailings, highlighting the benefit of deeper genomic and functional analyses to understand their ecological roles and evaluate their potential for heavy metal remediation.

## Conclusions

To our knowledge, this is the first study to apply integrated metagenomic and metaproteomic analyses to mine tailings microbial communities. Metagenomic sequencing revealed a broad range of novel taxa, particularly within the Actinobacteriota, Desulfobacterota, and uncultured phyla such as *Ca.* Eremiobacterota and *Ca.* SZUA-79, some of which were implicated in iron/sulfur cycling processes.

Functional profiling revealed distinct ecological strategies employed by differing microbial populations across geochemical gradients. In shallow, oxidized zones, iron- and sulfur-oxidizing Gammaproteobacteria were prevalent and active. Deeper, anoxic layers were dominated by potential sulfate-reducing organisms, many of which were taxonomically novel. Metal resistance mechanisms within communities as well as in individual taxa were highly diverse, reflecting potential adaptive diversity present within tailings microbiomes. The relative toxicity of metals was an important factor influencing MRG abundances in metagenomes. There were no clear correlations between MRG abundances and environmental metal concentrations, although organisms inhabiting highly contaminated tailings expressed similar MRGs targeting copper, nickel, and iron.

Although proteomics data was sparse and uneven from this challenging matrix, the expression of genes related to metal resistance and iron/sulfur cycling confirmed that identified metal-interacting microbes in these tailings not only encode, but actively utilize, a diverse set of MRGs and MMGs.

However, the low numbers of identified proteins in many samples restricted our ability to interpret the data in a systematic or quantitative manner.

The findings presented in this study highlight the need to develop improved methodologies for extracting nucleic acids and proteins from challenging environmental matrices like mine tailings.

However, despite substantial technical barriers towards obtaining sufficient DNA/protein for sequencing, including low biomass and interference from contaminants such as PEG, our results demonstrate the potential of multi-omics approaches to reveal new insights into microbial adaptations and activities within metal-contaminated environments.

## Methods

### Sampling and DNA extraction

Core sample collection and DNA extraction from mine tailings were performed as described in (Chen et al., 2024). Briefly, cores measuring 5.0–7.6 cm (2–3 inches) in diameter and 3–5 m in length (depth below surface level) were extracted using a drill and aluminum pipe, capped, and transported to the lab for storage at -20 °C prior to processing. Cores were sliced into 10 cm subsections, and 10 g of tailings from each subsample was weighed into a 50 mL falcon tube for DNA extraction using QIAGEN DNEasy PowerMax Soil kit. DNA samples were concentrated to a volume of 50-100 μL using ethanol/sodium acetate precipitation, and final DNA concentrations were determined using the Thermo Fisher Qubit dsDNA High Sensitivity Assay Kit. Samples were stored at -20 □ prior to sequencing.

### Geochemical assessments

Pore-water samplers (Soil Moisture Inc. model 1900 soil water solution samplers) were installed at each location at 10 to 20 cm depth intervals at ML25 and ML34 sampling locations, and water was collected from samplers after approximately one week. Piezometers were used to collect pore-water at NR3 (0.15–0.4 m intervals), as well as at NR18 below depths of 1.13 m. Pore-water pH, Eh, alkalinity, and electrical conductivity were measured on-site. Remaining water samples were filtered (0.45 µm) into 20 mL bottles and stored at 4 °C for further analysis: inductively coupled plasma-optical emission spectrometry and inductively coupled plasma-mass spectrometry (ICP-OES/ICP-MS) for cation concentrations, ion chromatography (IC) for anion concentrations, and non-dispersive infrared sensor (NDIR) analysis for total dissolved organic carbon (DOC).

### Metagenomic sequencing, assembly, and binning

A total of 112 DNA samples were obtained. Of these, 43 samples were sent for metagenomic sequencing based on 16S rRNA sequencing results (Chen et al., 2024). Selection criteria aimed to maximize the range of geochemistry and microbial community compositions observed across samples, while minimizing redundancy. For example, if a group of adjacent samples in one core had similar concentrations of dissolved metals, pH, and relative taxon abundances, one representative sample from the group was selected for metagenomic sequencing.

Paired-end metagenomic sequencing was performed at The Center for Applied Genomics (TCAG) (Toronto, Ontario), with an Illumina NovaSeq SP flowcell (2x150bp). Libraries were PCR-amplified prior to sequencing. Raw reads were decontaminated with BBduk (Bushnell, 2014), trimmed/quality filtered using sickle v1.33 (Joshi & Fass, 2011). SPAdes v3.15.5 (Bankevich et al., 2012) was used to assemble reads, with the --meta flag for metagenomes.

For binning, a set of assembled scaffolds ≥2,500 bp were filtered using pullseq v1.0.2 (Thomas, 2015). Bowtie2 v2.3.4.1 (Langmead & Salzberg, 2012) was used for read mapping based on sequencing depth to generate scaffold coverage information. Scaffolds were independently binned with three binning algorithms: CONCOCT (Alneberg et al., 2014), MetaBAT 2 (Kang et al., 2019), and MaxBin 2.0 (Wu et al., 2016), after which the dereplication, aggregation, and scoring tool (DAS Tool) (Sieber et al., 2018) was used to select the best set of high-quality non-redundant bins from the candidate bins. CheckM (Parks et al., 2015) was used to filter high-quality bins based on >70% completeness and <10% contamination thresholds.

After binning, MAGs were dereplicated across all samples with dRep (Olm et al., 2017) to generate a non-redundant set of MAGs for gene annotation.

### Metagenome annotation and data analysis

MAG taxonomy was assigned with GTDB-tk (v2.1.0, database release version 207.0) (Chaumeil et al., 2020). Phylogenetic trees were constructed with GToTree (Lee, 2019), and visualized with the online interactive tree of life (iToL) tool (Letunic & Bork, 2021). Iron-related genes were annotated with FeGenie (Garber et al., 2020), and sulfur metabolism genes were taken from DRAM (Distilled and Refined Annotation of Metabolism) annotations (Shaffer et al., 2020). DiSCo (Neukirchen & Sousa, 2021) was also used for sulfur gene annotation, which can distinguish between oxidative and reductive variants of *dsr* genes.

MRG gene annotation was based on the BacMet (v2.0) antibacterial biocide- and metal-resistance genes database (Pal et al., 2014). The BacMet 2.0 database contains 753 curated, experimentally verified bacterial metal and biocide resistance genes, as well as 155,512 predicted genes based on sequence similarity to the experimentally verified database (Pal et al., 2014). Initially, MRGs were annotated with the BacMet-scan software provided with the database, which uses a Basic Local Alignment Search Tool (BLAST) algorithm to identify similar genes. Prodigal (Hyatt et al., 2010) was used to predict open reading frames (ORFs) from MAGs for BacMet-scan, and results were filtered using a E<0.05 threshold.

The BacMet-scan method resulted in limited hits, so we developed an alternative method of MRG identification that would be comparable to the other gene annotation tools used in this study (FeGenie, DRAM, DiSCo) based on creating profile HMMs for MRGs of interest. Multiple sequence alignments (MSAs) were generated with MUSCLE (v3.8.31) (Edgar, 2004) for protein sequences associated with each of the 613 unique genes present in the combined BacMet experimentally verified and predicted gene databases. MSAs were verified by visual inspection in Geneious (v5.6) (Kearse et al., 2012). Profile HMMs were generated from MSAs using HMMER (v3.2.1) (Eddy, 2018), excluding genes with <5 representative sequences, resulting in 529 HMMs. The set of predicted ORFs used in the initial BacMet-scan were screened with the developed BacMet HMMs using HMMER, with E<0.05 indicating a significant hit.

To track the distribution of MRGs in MAGs compared to unbinned scaffolds, including mobile genetic elements, BacMet HMM hits for all scaffolds ≥2.5 kbp were aggregated and mapped to each scaffold’s classification from PPRmeta (Fang et al., 2019).

All data from FeGenie, DRAM, DiSCo, BacMet-scan, and BacMet HMM searches were exported to Python for analysis using the pandas (McKinney, 2010) library. Gene frequencies were aggregated into categories (*e.g.,* iron oxidation, sulfur reduction, copper resistance) and normalized to MAG relative abundance as a proxy for relative gene abundance, which was calculated from the average scaffold coverage of each MAG divided by the total coverage of all binned scaffolds within a sample. Figures were generated using matplotlib (Hunter, 2007).

### Protein extraction and sequencing

The full set of samples selected for metagenomics were also subjected to metaproteomic sequencing. The method of protein extraction was modified from a protocol described in Keiblinger et al. (2012), which compared the effectiveness of multiple protein extraction protocols on soil samples and found that an SDS-phenol extraction method (Wang et al., 2006) resulted in the highest numbers of mapped proteins from LC–MS/MS (Keiblinger et al., 2012). 10 g of tailings were measured into 50 mL falcon tubes and pre-washed with 10 mM Tris-EDTA buffer. The following was then added to sample tubes: 5 mL each of stainless steel and garnet beads, 15 mL of 1% SDS in 50 mM Tris-HCl pH 8.0, and 15 mL of Tris-HCl equilibrated liquid phenol. Samples were vortexed for 10 min and sonicated with a probe sonicator for 30 s, with all samples stored on ice between steps. Tubes were then placed onto a horizontal shaker at 160 rpm for 1 hr at room temperature, after which the vortex and sonication steps were repeated. Samples were centrifuged for 20 min at 3500 x g at 4 °C to separate the phenol layer (containing the solubilized proteins) from the aqueous layer. The phenol phase was transferred into a new 50 mL tube for precipitation of proteins overnight in 5 volumes of 0.1 M ammonium acetate in methanol at -20 °C.

To prepare peptides for LC-MS/MS, proteins were centrifuged for 20 minutes at 10,000 x g at 4 °C, washed with 2 mL of cold acetone, and resolubilized in 300 µL of denaturation buffer (8M urea in 50 mM Tris-HCl). Samples were then incubated with 10 mM dithiothreitol (DTT) at 60 °C for 30 mins to reduce disulfide bonds, then with 20 mM iodoacetamide (IAA) at room temperature for 30 mins to alkylate cysteines. Proteins were precipitated again in acetone, dried, and resuspended in 50 mM ammonium bicarbonate for overnight digestion with 1.5 µg trypsin at 37 °C. The digest was acidified with 1% formic acid, after which peptides were purified using Pierce C18 spin columns (Thermo Fisher #89870), dried in a vacuum centrifuge, and stored at -80 °C prior to LC–MS/MS.

LC-MS/MS was performed in the Biomolecular Core (BioCORE) Facility at Western University Schulich School of Medicine and Dentistry, London, Ontario, Canada, with the following specifications: peptides were resuspended in 1% acetonitrile (ACN) and 0.1% formic acid (FA), placed in an ultrasonic bath for 20 min, and centrifuged at 21,000 x g for 7 min at room temperature. Peptide samples were injected onto a Waters Acquity Ultra-Performance Liquid Chromatography (UPLC) M-Class system (Waters, Milford, MA) coupled to a Q Exactive Plus hybrid quadrupole-Orbitrap mass spectrometer (Thermo Fisher Scientific, Waltham, MA). Mobile Phase A consisted of water/0.1% FA and Mobile Phase B consisted of ACN/0.1% FA. Samples were trapped for 5 min at a flow rate of 5 μL/minute using 99% Buffer A, 1% Buffer B on a Symmetry C18 Trap Column, 5 μm, 180 μm × 20 mm (Waters).

Peptides were separated using a Peptide BEH C18 Column, 130Å, 1.7 μm, 75 μm × 250 mm operating at a flow rate of 300 nL/minute at 35 °C (Waters). Samples were separated using a nonlinear gradient consisting of 1–7% B over 1 minute, 7–23% B over 135 minutes and 23–35% B over 180 minutes before increasing to 99% B, for a total time of 3 h. The Q Exactive Plus mass spectrometer was controlled by Thermo Xcalibur software (v4.0). Data were acquired in the data-dependent acquisition mode (DDA) using a Fourier Transform/Higher-Energy Collision Dissociation (FT/FT/HCD) Top 12 scheme. MS1 survey scans were performed from 375 to 1,500 m/z at a resolution of 70,000 with Automatic Gain Control (AGC) set to 3e6 and a maximum injection time of 250 ms. Multiply charged peptide ions were isolated using the quadrupole with an isolation window of 1.2 m/z and were fragmented using HCD with Normalized Collision Energy (NCE) set to 25%. MS2 scans were performed at a resolution of 17,500 with AGC set to 2e5 and a maximum injection time of 64 ms. Dynamic exclusion was set to 40 s, and lock mass ion was enabled at 445.120025 m/z.

### Metaproteome annotation and data analysis

Initial data processing was conducted by personnel at the BioCORE facility, using the PEAKS 12 software (Bioinformatics Solutions Inc., Waterloo, Canada). To map predicted proteins, a custom ORF database was generated from the metagenomic sequencing data from this study by using Prodigal to predict ORFs in assembled scaffolds above 1 kbp in length. Predicted proteins from all metagenomes were combined into a single database to search the MS data against, with the following specifications: a precursor ion mass tolerance was set to 10 ppm, and a fragment ion mass tolerance was set to 0.02 Da.

Specific cleavage with trypsin was selected with a maximum of 2 missed cleavages. A fixed modification of carbamidomethylation and variable modifications of deamidation, oxidation and acetylation were added, and a maximum of two variable modifications were allowed per peptide. The peptide false discovery rate (FDR) was set to 1% for the database search. One unique peptide and an FDR of less than 1% were applied for confident protein identification.

Comparison of protein abundances across samples was not possible as all extracted peptides from each sample were injected to maximize the signal detection potential for low biomass samples, and therefore the amount of peptide injected varied between samples. Relative abundances within a sample were also not compared as the method of data collection did not account for differences in peptide detection ability by mass spectrometry. As a result, analysis of proteomic data was limited to inferring microbial activity based on the positive identification of associated proteins.

## Acknowledgements

We acknowledge the support of our industry partners at Glencore Sudbury INO for providing site access, historic site information, and field work assistance. We also thank the following members of Dr. David Blowes’ and Dr. Carol Ptacek’s research group for their help during field sampling: Mr. Austin Miller, Ms. Lisa Kester, Mr. YiZhi Yuan, and Mr. Luke Schofield. We thank Ms. Emilie Spasov for her help in tailings core processing and Ms. Rosalind Wang for her help with DNA extractions.

This work was supported by an Ontario Research Fund – Research Excellence grant (ORF-RE09-061). M.C. was supported by an NSERC CGS-D fellowship. D.S.G. was supported by an NSERC Banting postdoctoral fellowship. L.A.H. and D.W.B. were supported by the Canada Research Chairs.

Research reported in this publication was also supported by funding from Schulich School of Medicine and Dentistry, Canada Foundation for Innovation, Ontario Research Fund, and NSERC Research Tools and Instruments Awards to Western University through use of the Biomolecular Core (BioCORE) Facility.

## Author Contributions

Chen, Molly: Conceptualization, Methodology, Software, Formal analysis, Investigation, Data Curation, Writing – Original Draft, Writing – Review & Editing, Visualization

Hug, Laura A.: Conceptualization, Methodology, Resources, Writing – Review & Editing, Supervision, Project administration, Funding acquisition

Blowes, David W.: Resources, Writing – Review & Editing, Supervision, Project administration, Funding acquisition

Bain, Jeffrey G.: Investigation, Resources, Writing – Review & Editing, Project administration Grégoire, Daniel S.: Methodology, Writing – Review & Editing

## Data Availability

All molecular data associated with this project has been made publicly available. Metagenome sequence information is housed on NCBI under BioProject PRJNA1017420. Metagenome reads are available under SRA accessions SRR36271342- SRR36271385. MAGs are [currently submitted to NCBI and undergoing processing. Significant delays to this pipeline have existed for some years now – in the interim and for review purposes, the MAG sequence information can be found on the Open Science Framework at https://osf.io/v427n/overview?view_only=dc71d1b6927e4fd18e422b7746e236ab].

All proteomic data, including working files and raw data are available at the Open Science Framework at https://osf.io/v427n/overview?view_only=dc71d1b6927e4fd18e422b7746e236ab.

