## Supplemental figures and tables for "Diverse microbial metal resistance and novel metal cycling organisms in copper/nickel mine tailings"

### Supplementary Materials

Supplemental Tables S1 – S5 p. 2-4

Supplemental Figures S1 – S8 p. 5-12

**Table S1: Comparison of BacMet-scan method to custom HMMs for annotation of MRGs in protein sequences from MAGs**. The number and percentage of hits returned using each method are grouped based on metal. The second last row (Total) represents the sum of all MRG hits across all metals, which may include multiple hits for the same gene if it mapped to multiple categories. The Total (non-redundant, including non-metals) row represents the number of unique hits returned by either method, although this may still include multiple hits for the same query sequence to more than one unique gene in the BacMet database. Note that rare genes (<5 representative sequences in the BacMet database) were not used for the HMM method; instead, the BacMet-scan results for those genes (10,462 total, included in BacMet-scan totals in this table) were added to the HMM hits for data analyses.

| **Metal** | **BacMet-scan** | | **BacMet HMMs** | |
|---|---|---|---|---|
| | **Hits** | **% of Total** | **Hits** | **% of Total** |
| **Antimony (Sb)** | 7,687 | 2.18 | 26,031 | 1.69 |
| **Arsenic (As)** | 38,391 | 10.91 | 101,861 | 6.60 |
| **Bismuth (Bi)** | 841 | 0.24 | 2,297 | 0.15 |
| **Cadmium (Cd)** | 16,476 | 4.68 | 105,780 | 6.85 |
| **Chromium (Cr)** | 13,044 | 3.71 | 8,390 | 0.54 |
| **Cobalt (Co)** | 25,706 | 7.31 | 83,352 | 5.40 |
| **Copper (Cu)** | 45,067 | 12.81 | 209,372 | 13.56 |
| **Gallium (Ga)** | 6,856 | 1.95 | 25,150 | 1.63 |
| **Gold (Au)** | 911 | 0.26 | 10,951 | 0.71 |
| **Iron (Fe)** | 19,526 | 5.55 | 179,110 | 11.60 |
| **Lead (Pb)** | 8,084 | 2.30 | 23,364 | 1.51 |
| **Magnesium (Mg)** | 6,538 | 1.86 | 8,552 | 0.55 |
| **Manganese (Mn)** | 9,140 | 2.60 | 75,419 | 4.89 |
| **Mercury (Hg)** | 19,350 | 5.50 | 66,106 | 4.28 |
| **Molybdenum (Mo)** | 17,462 | 4.96 | 59,265 | 3.84 |
| **Nickel (Ni)** | 30,703 | 8.73 | 117,354 | 7.60 |
| **Selenium (Se)** | 6,973 | 1.98 | 3,995 | 0.26 |
| **Silver (Ag)** | 6,576 | 1.87 | 81,469 | 5.28 |
| **Tellurium (Te)** | 10,962 | 3.12 | 7,355 | 0.48 |
| **Tungsten (W)** | 27,111 | 7.71 | 114,477 | 7.42 |
| **Vanadium (V)** | 968 | 0.28 | 3,080 | 0.20 |
| **Zinc (Zn)** | 33,436 | 9.50 | 230,821 | 14.95 |
| **Total** | **351,808** | | **1,543,551** | |
| **Total (non-redundant, including non-metals)** | **329,520** | | **1,660,455** | |

**Supplemental Data File 1:**

**Table S2: Pore-water geochemistry measurements.** Adapted from Chen et al., 2024. Values for pH, Eh, alkalinity, electrical conductivity (EC), dissolved organic carbon (DOC), and total sulfur are provided along with the concentrations of select anions and cations for water samples taken at different depths at each sampling location (ML25, ML34, NR18, NR3). Cation/anion concentrations are provided if measured values exceeded detection limits for >50% of samples within a location.

**Table S3: Metal resistance gene counts by metal, sample, and scaffold origin.** Hits for MRGs are grouped by the targetted metal (columns), and by sample (rows). Within samples, MRGs are further divided into those on binned scaffolds, unbinned chromosomal scaffolds, unbinned plasmids, and unbinned phages. Only scaffolds >2.5 kbp in length were included in this analysis. Gene annotations were based on custom HMMs generated from metal resistance genes in the BacMet database (Pal et al., 2014).

**Table S4: Metal resistance gene abundances by metal, sample, and scaffold origin.** Hits for MRGs are grouped by the targetted metal (columns), and by sample (rows), and normalized to the percent relative abundances of the associated scaffold within each respective sample to calculate the relative gene abundance. Within samples, MRG abundances are further divided into those on binned scaffolds, unbinned chromosomal scaffolds, unbinned plasmids, and unbinned phages. Only scaffolds >2.5 kbp in length were included in this analysis. Gene annotations were based on custom HMMs generated from metal resistance genes in the BacMet database (Pal et al., 2014).

**Table S5: Summary of iron-related protein families that are represented as profile HMMs in FeGenie.** Adapted from Table 1 in Garber et al., (2020); see the associated publication for more information, including references to publications and databases for protein families.

| Function | Protein Families |
|---|---|
| **Iron transport** | EfeUOB, FbpABC, SfuABC, YfuABC, FeoAB(C), FutA1, FutA2, FutB, FutC, YfeABCD |
| **Heme oxygenase** | ChuS, ChuZ, HemO, PigA, HemRSTUV, HmoB, HmuO, HugZ, HupZ, Isd-LmHde, IsdG, IsdI, MhuD, PhuS (in PhuRSTUVW) |
| **Heme transport** | HasRADE(B)F, HmuRSTUV, HmuY, HmuY', HutZ, HxuCBA, IsdX1, IsdX2,PhuRSTUVW, Rv0203 |
| **Transferrin/Lactoferrin** | TbpAB (LbpAB), SstABCD |
| **Siderophore synthesis** | AcsABCDEF, AmoA, AngR, AsbABCDEF, DhbACEBF, entD-fepA-fes-entF-fepECGDB-entCEBA-ybdA, IroD in IroNBCDE, IucABCD,, IutA, MbtIJABCDEFGH, LbtA (in LbtUABC), PchABCDREFGHI, PvdQAPMNOFEDJIHLGS, PvsABCDE, VenB, Vab genes in VabR-fur-vabGA-fur-VabCEBSFH-fur-fvtA-vabD, Vib genes in VibB-vibEC-vibA-vibH-viuPDGC-vibD and viuAB-vibF, RhbABCDEF-rhrA-rhtA |
| **Siderophore transport** | BesA, CbrABCD, TonB-ExbB-ExbD, FatABCD, FecIRABCDE, FeuABC-yusV, FhuACDB, FhuF, FptABCX, FpuAB, FpuC, FpuD, FpvIR-FpvA-FpvGHJKCDEF, FvtA in VabR-fur-vabGA-fur-VabCEBSFH-fur-fvtA-vabD, HatCDB, IroNBCDE, LbtUABC, PirA, PiuA, PvuABCDE, Viu genes in VibB-vibEC-vibA-vibH-viuPDGC-vibD and viuAB-vibF, YfiZ-yfhA, YfiY, YqjH, ybdA and Fep genes in entD-fepA-fes-entF-fepECGDB-entCEBA-ybdA |
| **Iron gene regulation** | DtxR63, FecR (in FecIRABCDE), FeoC in FeoAB(C), Fur, IdeR, YqjI, RhrA in RhbABCDEF-rhrA-rhtA |
| **Iron oxidation** | Cyc1, Cyc2 (cluster 1), Cyc2 (cluster 2), FoxABC, FoxEYZ, Sulfocyanin, PioABC |
| **Iron reduction** | CymA, MtrCAB, OmcF, OmcS, OmcZ, FmnA-dmkA-fmnB-pplA-ndh2-eetAB-dmkB, DFE_0448-0451, DFE_0461-0465 |
| **Possible iron oxidation /reduction** | MtoAB, Cyc2 (cluster 3) |
| **Probable iron reduction** | MtrCB, MtrAB, MtoAB-MtrC |
| **Iron storage** | Bfr, DpsA, Ftn |
| **Magnetosome formation** | MamABEKLMOPQI (Note: These genes are found in all known magnetotactic microorganisms, except for *mamL* which is found in magnetite-producing magnetotactic microorganisms) |

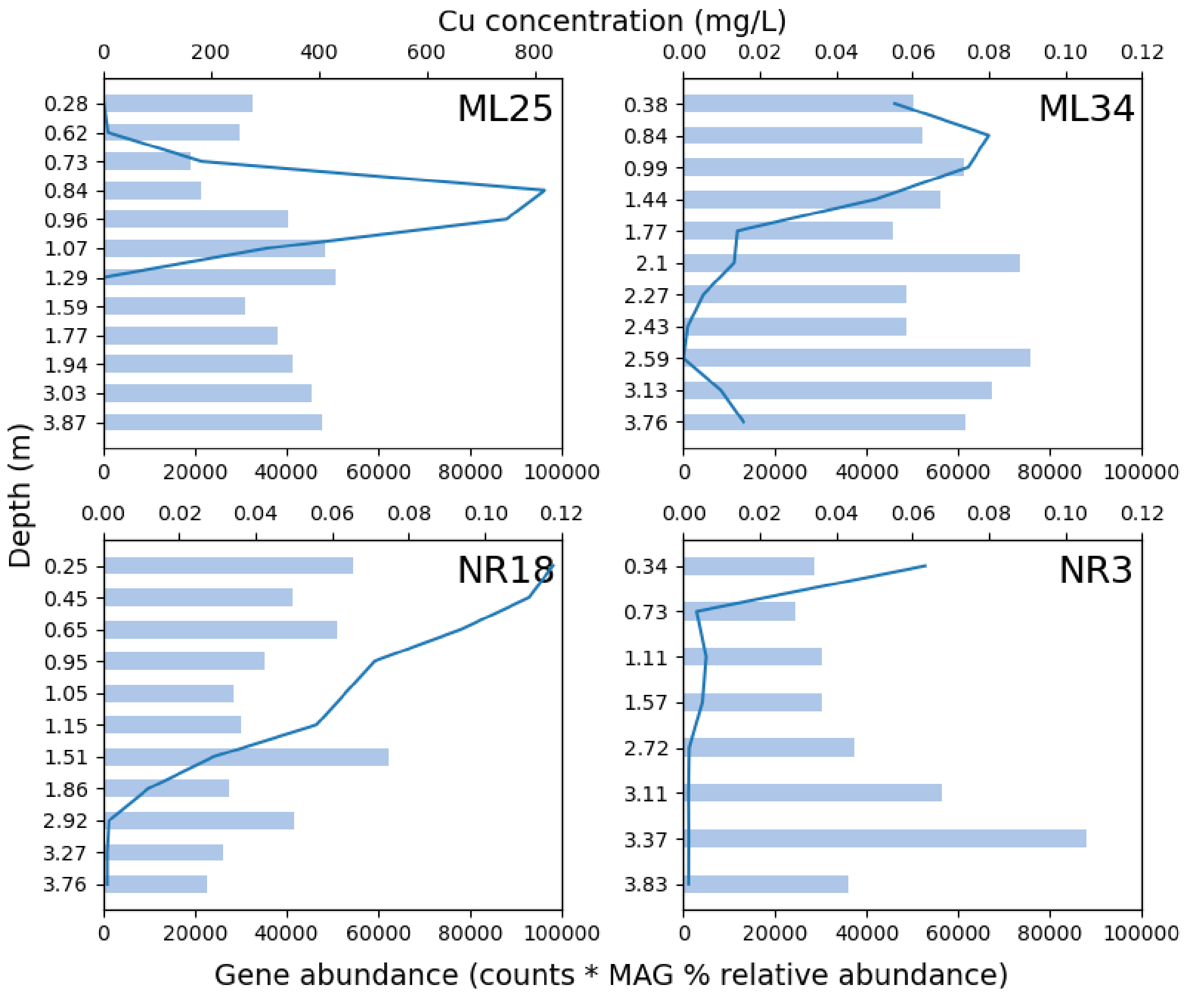


**Figure S1: Abundance of copper resistance genes and copper concentrations along four vertical mine tailings cores**. Bar plots (lower x-axis values) represent relative gene abundances for Cu-associated MRGs. Line plots (upper x-axis values) represent Cu concentrations in tailings pore-water. Metal concentrations in ML25 are reported on a different scale for visibility. There was no significant correlation between copper concentrations and MRG abundances across all samples ($p=0.18$) or across samples within each tailings core ($0.15 \leq p \leq 0.51$). Note that the y axes are not scaled proportionally to depth.

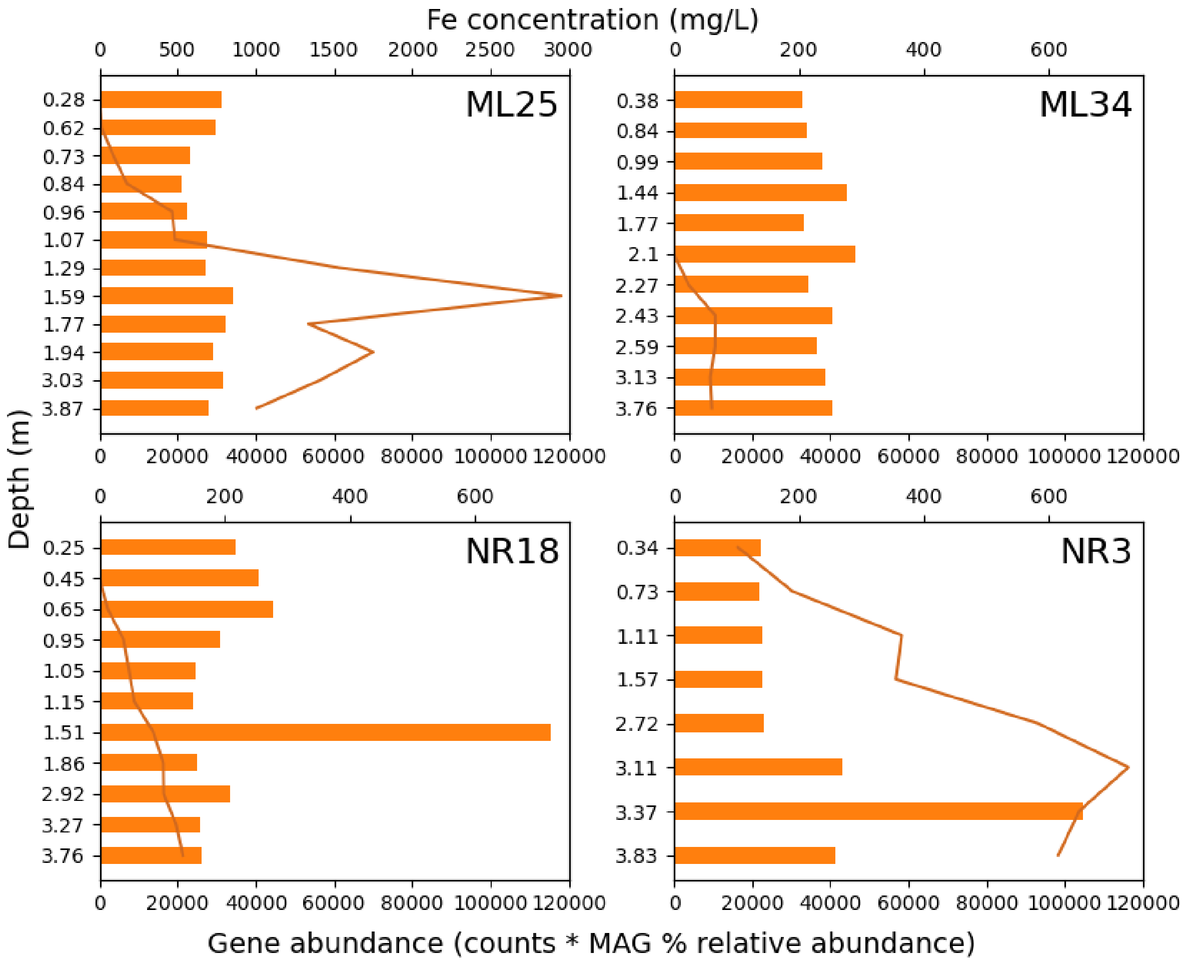


**Figure S2: Abundance of iron resistance genes and iron concentrations along four vertical mine tailings cores**. Bar plots (lower x-axis values) represent relative gene abundances for Fe-associated MRGs. Line plots (upper x-axis values) represent Fe concentrations in tailings pore-water. Metal concentrations in ML25 are reported on a different scale for visibility. There was no significant correlation between iron concentrations and MRG abundances across all samples ($p=0.79$) or across samples within each tailings core ($0.05 \leq p \leq 0.95$). Note that the y axes are not scaled proportionally to depth.

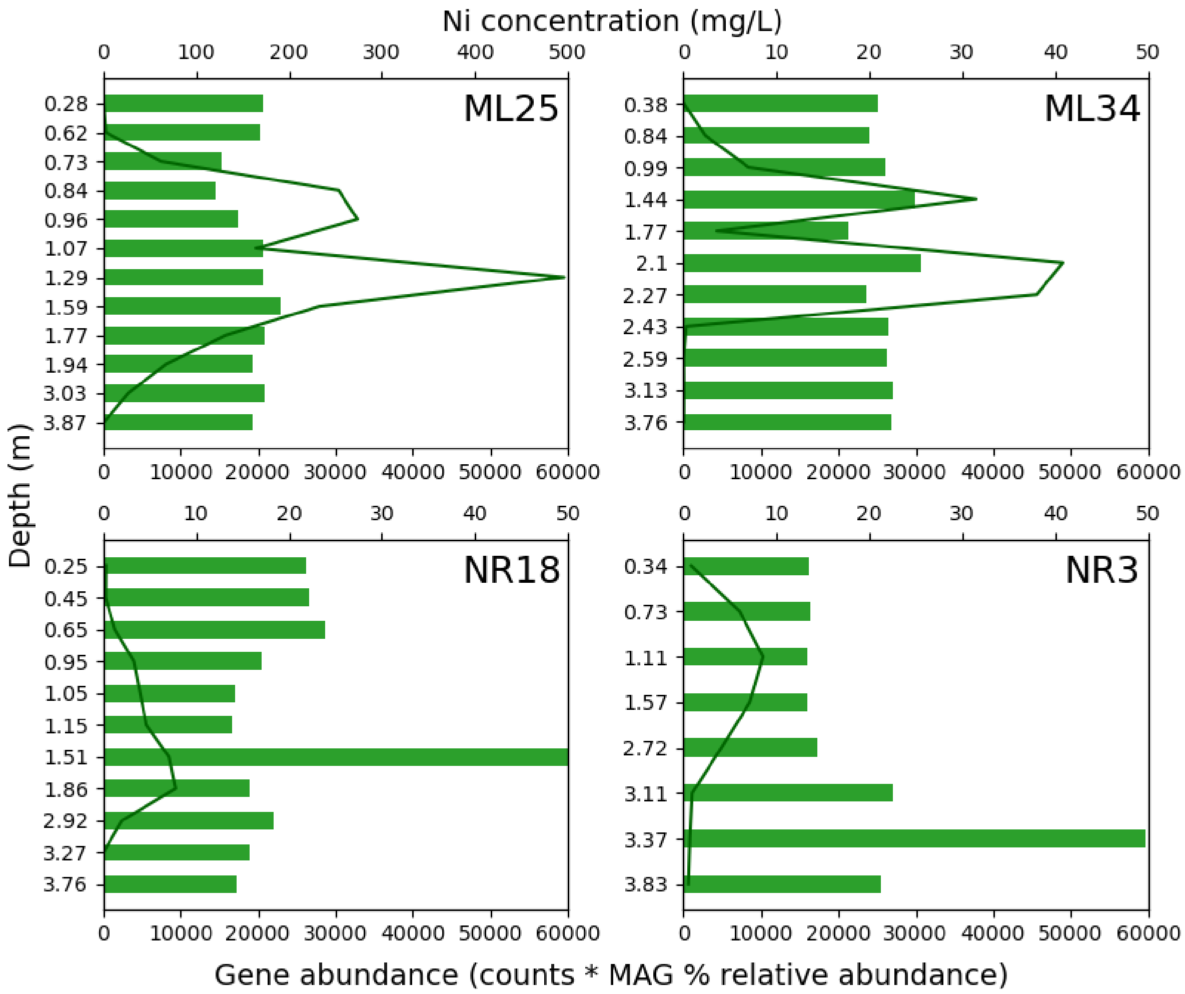


**Figure S3: Abundance of nickel resistance genes and nickel concentrations along four vertical mine tailings cores**. Bar plots (lower x-axis values) represent relative gene abundances for Ni-associated MRGs. Line plots (upper x-axis values) represent Ni concentrations in tailings pore-water. Metal concentrations in ML25 are reported on a different scale for visibility. There was no significant correlation between nickel concentrations and MRG abundances across all samples ($p=0.26$) or across samples within each tailings core ($0.17 \leq p \leq 0.92$). Note that the y axes are not scaled proportionally to depth.

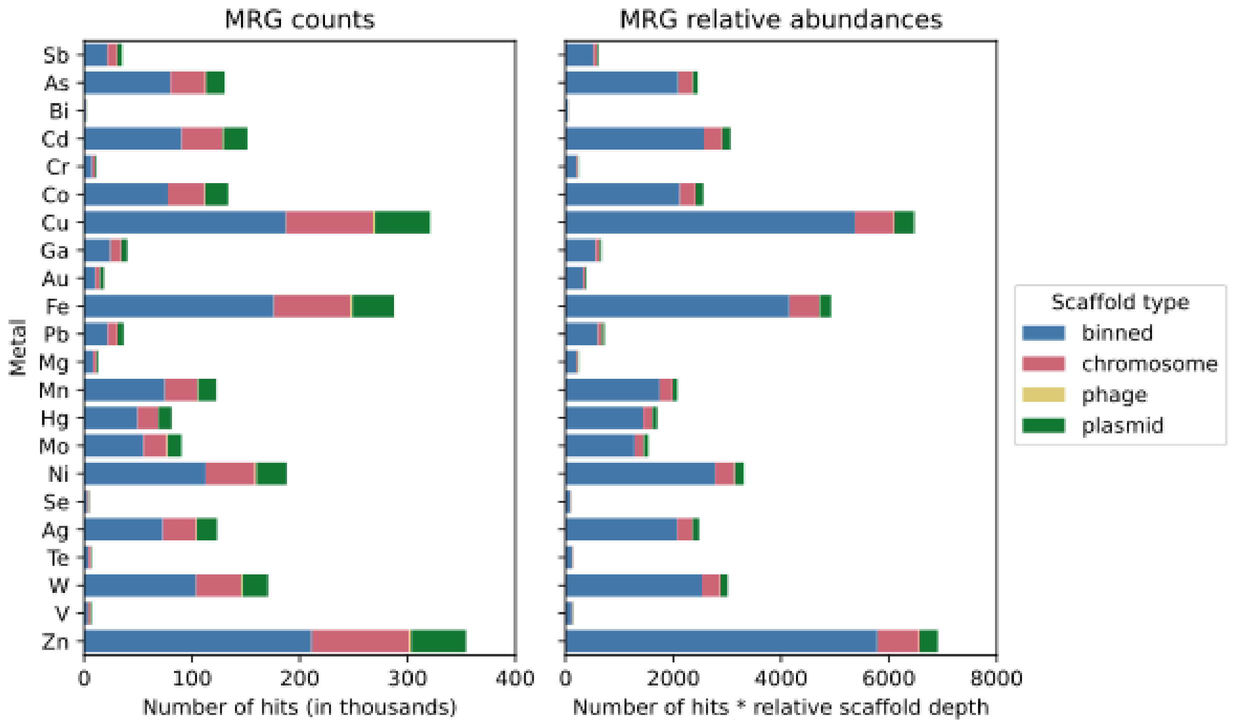


**Figure S4: Distribution of metal resistance genes in tailings across binned and unbinned scaffolds.** MRGs on all assembled scaffolds >2.5 kbp in length were annotated using HMMs of genes in the BacMet database. Rare genes (<5 representatives in the BacMet database) were annotated with BacMet-scan**.** Hits were agreggated based on metal(s) and displayed as either counts (left), or relative gene abundances (hits multiplied by relative scaffold abundances within their respective sample) (right). Bars are colored based on whether scaffolds are binned or unbinned, with unbinned scaffolds divided into chromosome (unbinned genomic DNA), phage, and plasmid categories, which were determined using PPRmeta.

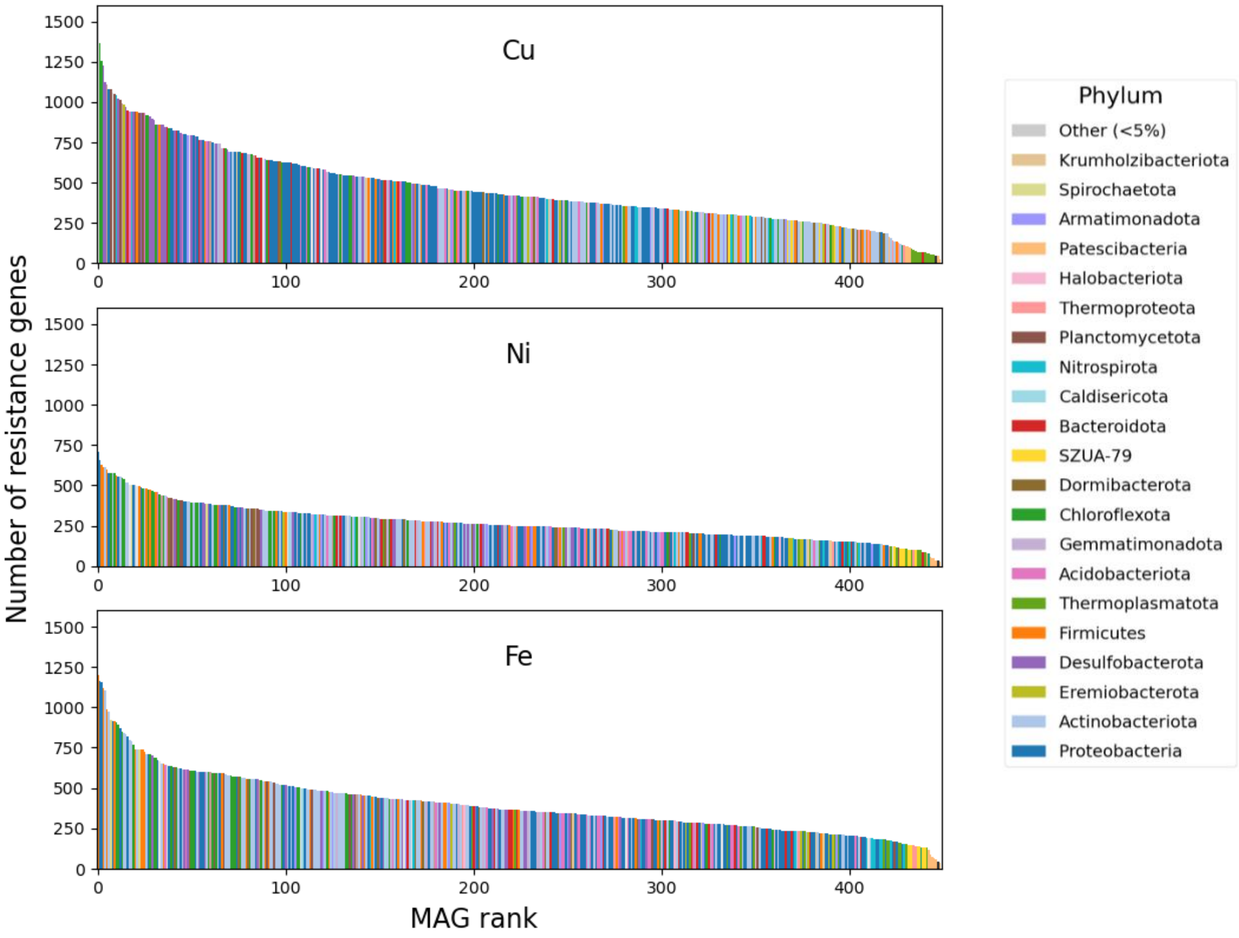


**Figure S5: MAG rank abundance plot by metal resistance genes**. Bars representing all non-redundant MAGs across all tailings samples are ordered from highest to lowest by counts of predicted MRGs targeting copper (top), nickel (middle), and iron (bottom). MRGs were annotated using profile HMMs based on the BacMet resistance gene database. Bars indicate the total number of MRGs encoded on a MAG, and are colored by the corresponding phylum (GTDB-tk v2.1.0) of each MAG.

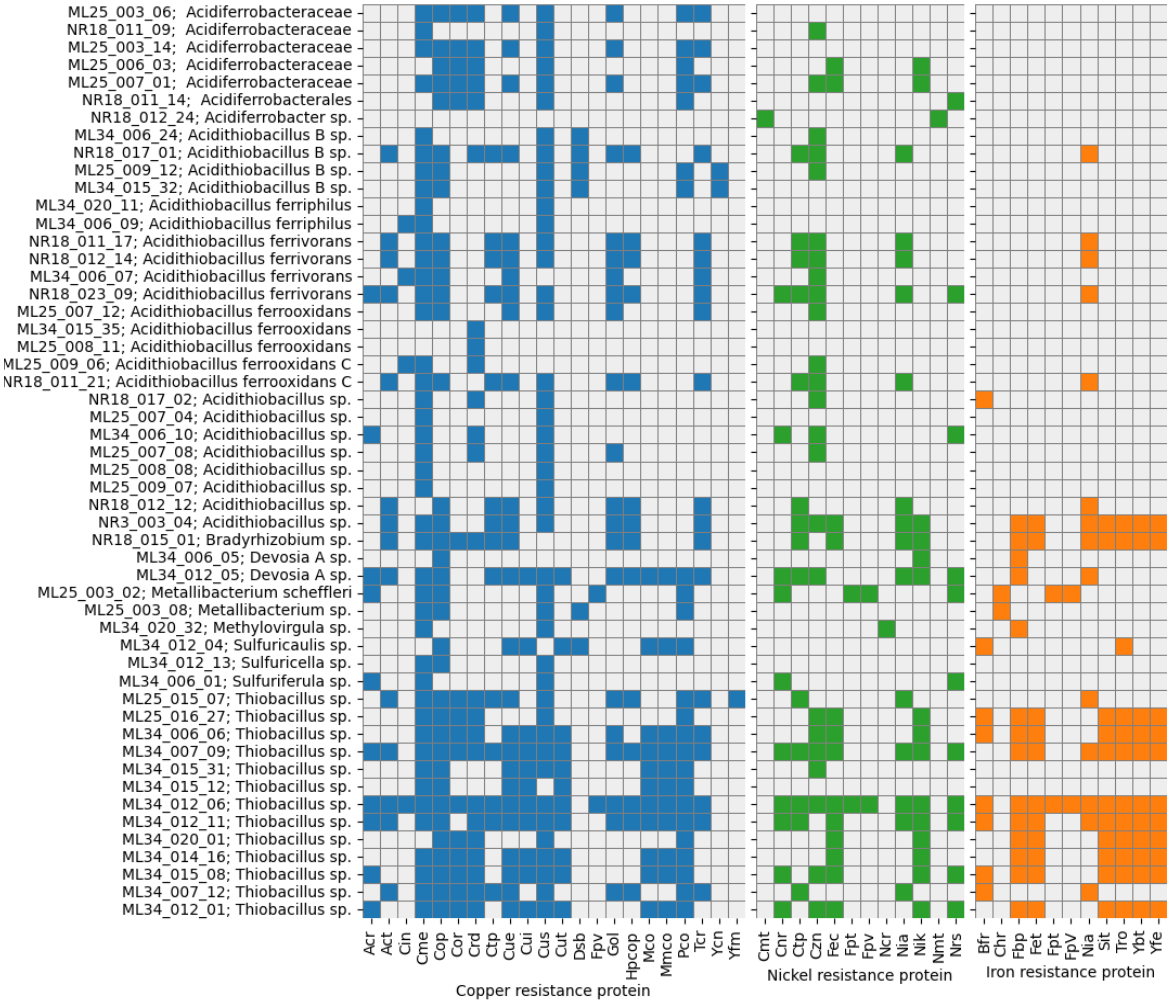


**Figure S6: Taxonomic distribution of copper, nickel, and iron resistance proteins in tailings communities**. Shaded cells represent positive detection of specific proteins associated with copper, nickel, and iron resistance. Proteins within common pathways (*e.g.,* Cop, Cus) were aggregated. Proteins were identified by mass spectrometry and mapped to predicted proteins from metagenomic sequencing, and aggregated by the MAG from which they originated (if any). Bins are labelled by their ID, followed by their lowest level of GTDB-assigned taxonomy. Only bins with >50 unique proteins detected across all samples were included in this analysis.

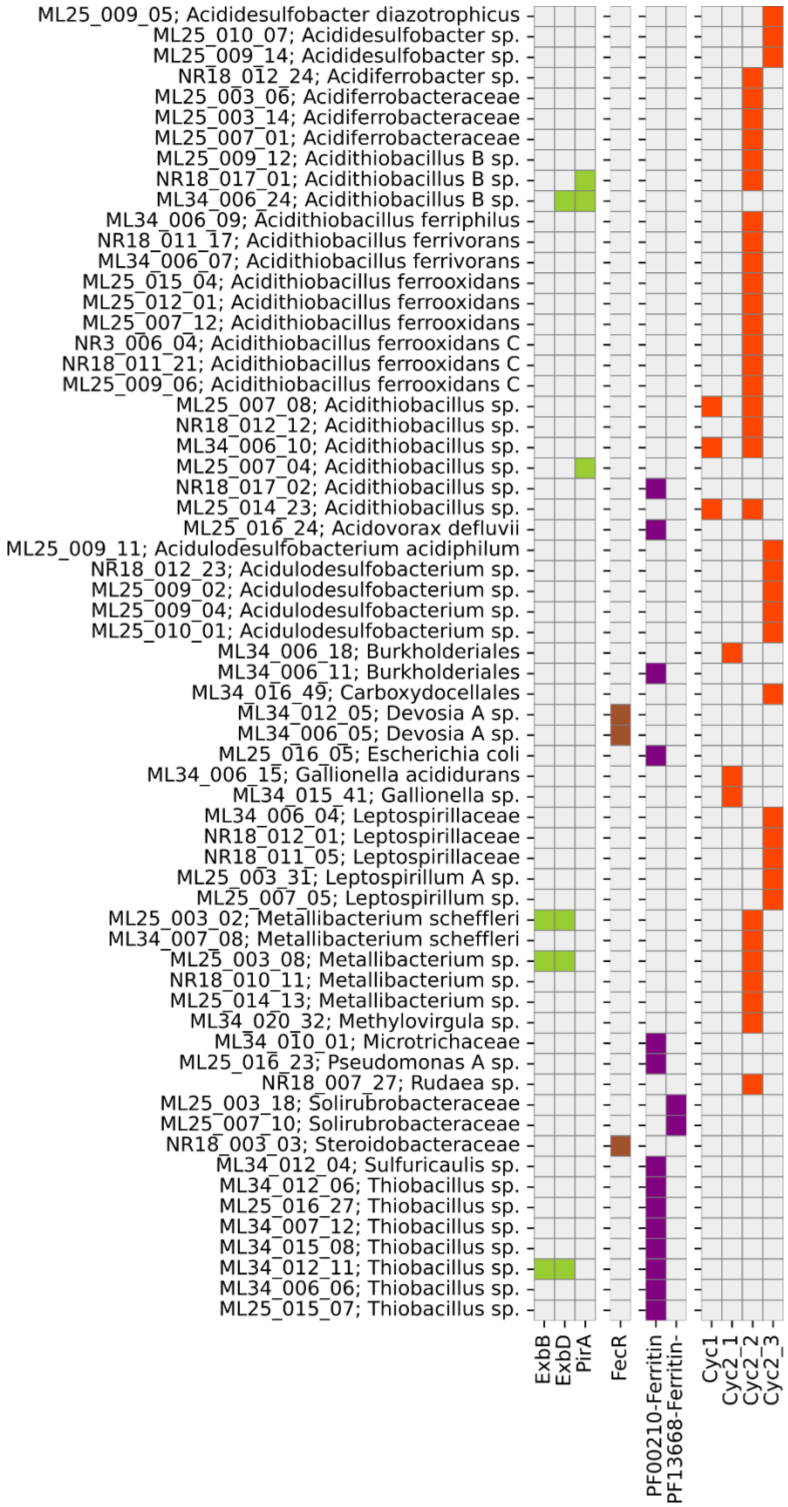


**Figure S7: Taxonomic distribution of iron-related proteins in tailings communities**. Shaded cells represent positive detection of specific proteins associated with (left to right): siderophore synthesis, iron gene regulation, iron storage, and iron oxidation. Proteins were identified by mass spectrometry and mapped to predicted proteins from metagenomic sequencing, and aggregated by the MAG from which they originated (if any). Bins are labelled by their ID, followed by their lowest classified level of GTDB-assigned taxonomy.

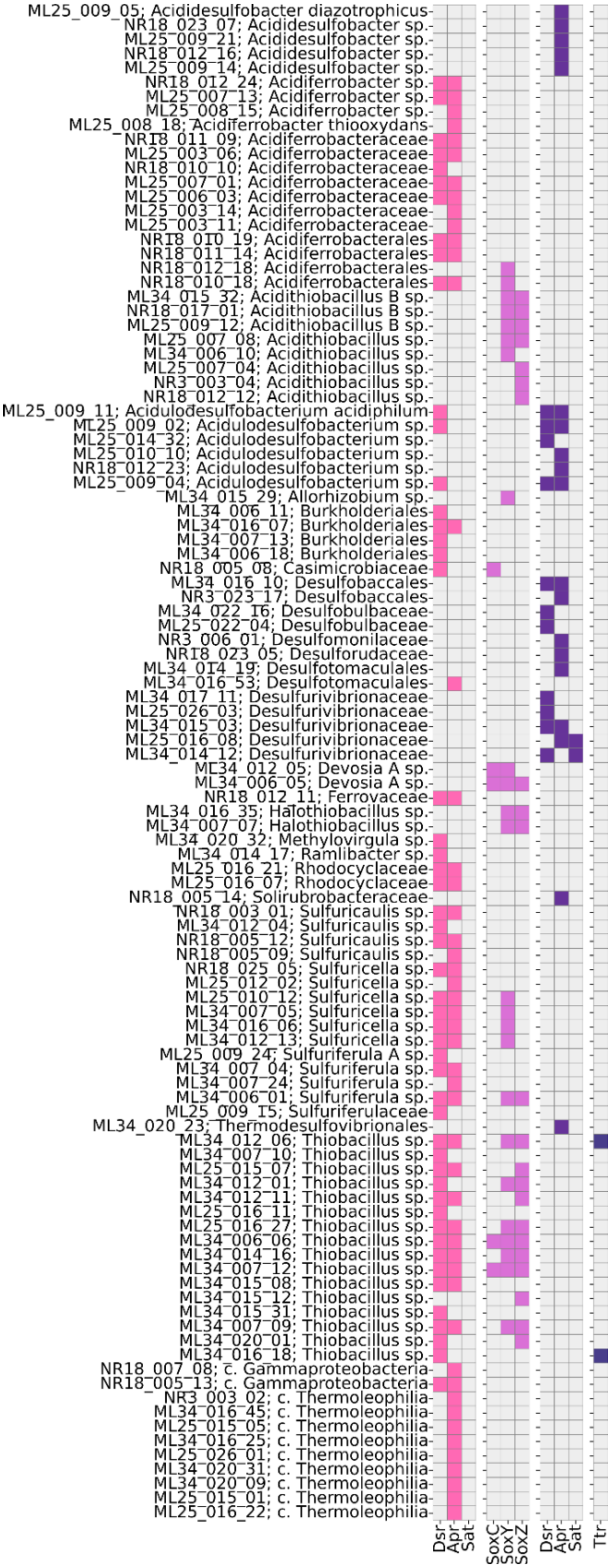


**Figure S8: Taxonomic distribution of sulfur-related proteins in tailings communities**. Shaded cells represent positive detection of specific proteins associated with (left to right): sulfur oxidation (Dsr pathway), sulfur oxidation (Sox pathway), dissimilatory sulfate reduction (Dsr pathway), and tetrathionate reduction (Ttr). Proteins were identified by mass spectrometry and mapped to predicted proteins from metagenomic sequencing, and aggregated by the MAG from which they originated (if any). Bins are labelled by their ID, followed by their lowest classified level of GTDB-assigned taxonomy.
